# CryoEM architecture of a native stretch-sensitive membrane microdomain

**DOI:** 10.1101/2023.08.25.554800

**Authors:** Jennifer M. Kefauver, Markku Hakala, Luoming Zou, Josephine Alba, Javier Espadas, Maria G. Tettamanti, Leandro F. Estrozi, Stefano Vanni, Aurélien Roux, Ambroise Desfosses, Robbie Loewith

## Abstract

Biological membranes are partitioned into functional zones containing specific lipids and proteins, termed membrane microdomains. Their composition and organization remain controversial owing to a paucity of techniques that can visualize lipids *in situ* without disrupting their native behavior^1,2^. The yeast eisosome, a membrane compartment scaffolded by the BAR-domain proteins Pil1 and Lsp1, senses and responds to mechanical stress by flattening and releasing sequestered factors^3–7^. Here, we isolated native eisosomes as helical filaments of Pil1/Lsp1 lattice bound to plasma membrane lipids and solved their structures by helical reconstruction. We observe remarkable organization within the lipid bilayer density from which we could assign headgroups of PI(4,5)P_2_ and phosphatidylserine bound to Pil1/Lsp1 and a pattern of membrane voids, signatures of sterols, beneath an amphipathic helix. We verified these assignments using *in vitro* reconstitutions and molecular dynamics simulations. 3D variability analysis of the native eisosomes revealed a dynamic stretching of the Pil1/Lsp1 lattice that affects functionally important lipid sequestration, supporting a mechanism in which membrane stretching liberates lipids otherwise anchored by the Pil1/Lsp1 coat. Our results provide mechanistic insight into how eisosome BAR-domain proteins create a mechanosensitive membrane microdomain and, more globally, resolve long-standing controversies about the architecture and nature of lipid microdomains.

## Introduction

Over fifty years ago, it was proposed that the plasma membrane is compartmentalized into laterally heterogenous domains^8,9^. Although the biological evidence for membrane compartmentalization is overwhelming (e.g. localization of immune receptors to detergent-resistant or liquid-ordered membrane domains^10–12^, spatiotemporal organization of transmembrane proteins and plasma membrane lipids by the cortical actin cytoskeleton^13–15^) the determinants and the physical structure of the lipid organization within the membrane remain controversial. This is because almost all tools used to study membrane lipids also risk perturbing their behavior within the membrane context^1,2,16^.

In *S. cerevisiae*, at least three membrane compartments have been identified: the membrane compartment containing Pma1 (MCP), the membrane compartment containing Can1 (MCC), and the highly dynamic membrane compartment containing TORC2 (MCT)^3^, in addition to the patchwork organization of many other integral membrane proteins^17^. The MCC microdomains, also known as eisosomes, are randomly distributed membrane furrows, about 300nm long and 50 nm deep, scaffolded by the Bin/amphiphysin/Rvs (BAR) domain family protein Pil1 and its paralog Lsp1^3,4,18^. The eisosomes are unique among membrane compartments in their stability; under steady-state conditions, they do not change size or distribution within the membrane for the duration of the cell cycle and they are non-overlapping with sites of endocytosis, protecting their resident proteins from turnover^19–21^. The function of eisosomes remains mysterious, but they are clearly implicated in sensing and responding to plasma membrane stress: various stimuli including hypo-osmotic shock, heat shock, and mechanical pressure cause eisosomes to flatten and release the sequestered eisosome-resident proteins, enabling their signaling and transport _functions3–5,18,22,23_.

Eisosomes are a plasma membrane feature that is unique to fungi and some algae^24^. However, they may have functional relatives in mammalian cells. They share many features with caveolae, including flattening in response to membrane stress^25,26^, their role in regulating plasma membrane homeostasis^27,28^ and assembly dependent on negatively-charged lipids (phosphatidylserine and PI(4,5)P_2_)^29–31^. There are also several examples of mammalian membrane domains scaffolded by BAR-domain proteins including the t-tubules of cardiac and skeletal muscle tissue that are stabilized by BIN-1^32–34^ and endosomal tubulation by SNX family proteins and pacsin-2^35^.

BAR domain proteins are a large and diverse family of proteins that play physiological roles in curvature sensing and/or induction at cellular membranes^33,36^. The majority have a characteristic banana-shape with a membrane-binding surface that displays dense positive charge, enabling the interaction with negatively-charged lipid headgroups^33,35–37^. They have been shown to induce curvature in membranes via the combination of a variety of mechanisms including 1) their own intrinsic curvature, 2) the insertion of amphipathic motifs into the membrane, and 3) their self-organization into large protein lattices^33,35^. While a wealth of data exists on the features of BAR domain proteins that mediate their curvature sensing/generation function, as well as their lipid binding affinities *in vitro*, what remains unknown is how BAR scaffolding impacts the organization of the complex mixture of lipids naturally found in cell membranes *in vivo*.

Here we have isolated an intact membrane microdomain, scaffolded in helical tubules by the eisosome BAR-domain proteins Pil1/Lsp1. By solving high resolution structures using cryoEM and helical reconstruction, we have a unique opportunity to visualize the unperturbed eisosome membrane lipids which exhibit a surprising level of organization. A pattern of small membrane voids beneath the amphipathic helix of Pil1/Lsp1 forms a helical striation in the protein-bound cytoplasmic leaflet, but not the exoplasmic leaflet, and lipid headgroup density is observed within charged pockets of the Pil1/Lsp1. Using a combination of *in vitro* reconstitution and molecular dynamics (MD) simulations with various lipid combinations of known composition, we were able to identify which lipid species are responsible for the structural signatures we found in the native eisosome membrane microdomain, as well as observe a reduction in the dynamics of all the lipid species present at the Pil1 microdomains. The membrane voids are footprints of sterol molecules stabilized at the sites of bulky sidechains of the amphipathic helix and the bound lipid density is consistent with PI(4,5)P_2_ and phosphatidylserine headgroups bound in each charged pocket. Remarkably, we also observe a dynamic stretching behavior of the Pil1/Lsp1 lattice in the native samples which alters the organization of these lipids in the cytoplasmic leaflet, revealing a possible mechanism of membrane tension sensing by the eisosomes.

### Isolation of native-like eisosome filaments

Eisosome tubules were isolated from *S. cerevisiae* expressing a TAP-tag on the Target of Rapamycin Complex 2 (TORC2) subunit Bit61. Using a gentle purification procedure involving hand-grinding of flash frozen yeast under liquid nitrogen and an initial extraction buffer containing CHAPS detergent at less than 1/10x critical micelle concentration (0.5mM CHAPS in extraction buffer versus CMC of ∼6mM at 250mM NaCl)^38^, we were able to preserve a lattice of Pil1/Lsp1 eisosome structural proteins in a near-native state, bound to the intact plasma membrane bilayer, observable as a two layer density within the protein tubule (Fig 1A). Pil1 is a common contaminant in *S. cerevisiae* pull-downs^39^ and the amount of Pil1/Lsp1 protein isolated by our methods is below the detection limit of our protein gels, suggesting our eisosome filaments are a contamination of our intended target (Fig Ext Data 1A). Nevertheless, the large tubulated structures they form were salient features on the EM grid, enabling us to collect a sufficiently large dataset for structural determination via helical reconstruction (Fig Ext Data 1B).

**Figure 1.**
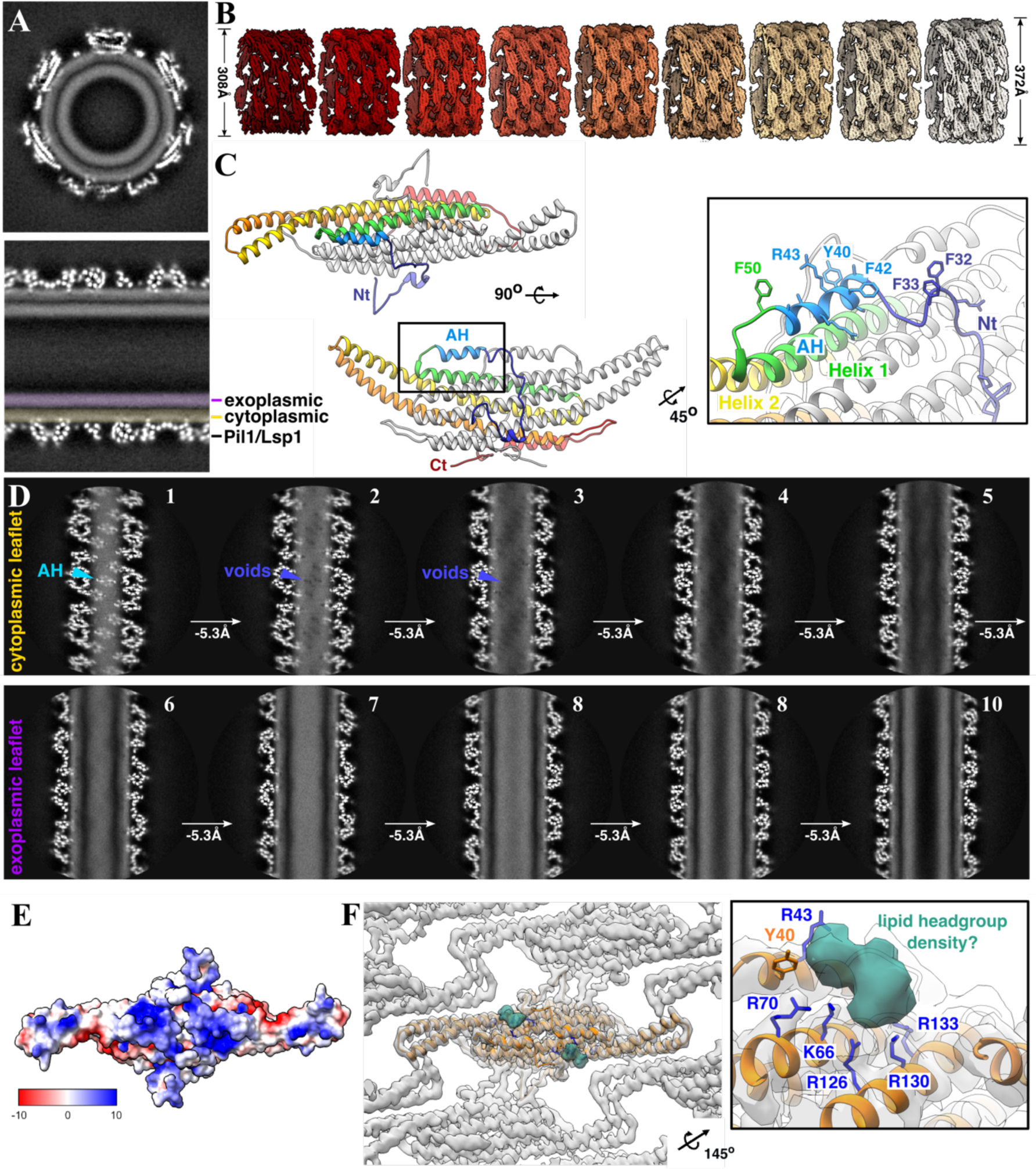
Native-like eisosomes retain unperturbed plasma membrane microdomain. **A.** Central (“transverse” above and “sagittal” below) slices of helical reconstruction of a native eisosome with membrane bilayer visible. **B.** Sharpened maps of 9 helical structures of varying diameter. **C.** Model of Pil1 dimer, rainbow coloring on chain A from Nt (blue) to Ct (red). [inset] Zoom on AH. **D.** Series of one-pixel slices through helical reconstruction of native eisosome, separated by ∼5.3Å depth. Cyan arrow indicates AH (panel 1), violet arrow indicates membrane voids (panels 2-3). Void pattern continues through cytoplasmic leaflet (top panels 1-5) but is absent in exoplasmic leaflet (bottom panels 6-10). **E.** Electrostatic surface prediction of Pil1 model. **F.** Unassigned putative lipid density (sea green) in deepEMhancer sharpened map localized to charged pocket. [inset] Charged residues coordinating unassigned density.

While eisosomes have been shown to have a furrow-like halfpipe structure *in vivo*^29,40^, our isolated eisosome tubules appear to be closed, continuous helices of Pil1/Lsp1 proteins (Fig 1A&B). Similar tubular structures have previously been shown in reconstituted Pil1 samples and eisosomes purified from yeast^29^. Presumably, once the eisosomes are freed from the plasma membrane by our gentle purification methods, the Pil1/Lsp1 lattice realigns to form a helical conformation around a lower-energy tubular state of the plasma membrane lipids to which it is bound. Alternatively, the eisosomes that we have isolated were already in a tubulated state *in vivo*, a behavior that has been observed in deletion strains of the eisosome-resident Sur7-family proteins, in palmitoylcarnitine-treated *S. cerevisiae*^41^, and upon Pil1 overexpression in *S. pombe*^42,43^.

In our raw images and 2D classes, we noted a large variation in tubule diameters (Fig Ext Data 1B&C). Using helical reconstruction, we were able to resolve 9 eisosome tubule structures with different helical symmetries and tubule diameters ranging from 308-372Å, but with a nearly identical lattice pattern of Pil1/Lsp1 dimers in all structures (Fig 1B, Ext Data 1D, Ext Data 2A, Supp Table 1&2). These tubule diameters are within the range of diameters of curvatures observed in eisosomes *in situ* (∼300-600Å)^44^.

### Structure of native Pil1/Lsp1 lattice

To improve resolution and enable 3D classification of the Pil1/Lsp1 dimers, we used a symmetry expansion and particle subtraction strategy, masking a central dimer and each of its contacting partners (a total of 7 dimers) combined with a small spherical mask to include the lipid bilayer beneath the central dimer (Fig Ext Data 2B). This allowed us to merge lattice pieces from the 9 helical structures (resolution ranging from 3.7Å-7.2Å) into an expanded dataset (Supp Table 2). Refinement of this dataset improved resolution to ∼3.2Å, enabling us to build and refine models of both Pil1 and Lsp1 proteins (Fig 1C, Ext Data 3A&B, Ext Data 4A).

Because our samples most likely contain a mixture of both Pil1 and Lsp1 proteins and their sequence conservation is very high, we could not conclusively differentiate between these two proteins in our structures (Fig Ext Data 3C, Ext Data 5, 73% identity, *MUSCLE*). Although there is some evidence to suggest that Pil1 and Lsp1 form heterodimers in the cytosol prior to their assembly at the plasma membrane^45^, it is also known that they are capable of homodimerizing and tubulating membrane *in vitro*^29^. While deletion of Lsp1 does not cause profound alteration in eisosome structure at the plasma membrane, deletion of Pil1 prevents the formation of eisosomes^40,46^. Due to its essential role, we have chosen to base our interpretations on the model of a Pil1 homodimer.

The overall “banana-shaped” BAR domain structure of the Pil1 dimer in our model corresponds well with the previous crystal structure of an Lsp1 dimer^47^ (Fig 1C). Each monomer contributes three antiparallel alpha-helices that form kinked coiled-coil domains, while a fourth alpha-helix from each monomer sits atop the convex surface of the curved dimer. We were able to observe two additional structured regions of the protein in our maps: 1) a folded N-terminus (Nt) that forms lattice contact sites with the Nt of a neighboring dimer followed by 2) an Nt amphipathic helix (AH) buried within the lipid density of the “inner leaflet” of the bilayer that runs parallel to the BAR domain helices of each Pil1/Lsp1 monomer (Fig 1C, inset).

Previous nanometer-resolution helical reconstructions of reconstituted Pil1 and Lsp1 proteins revealed a lattice pattern that could be fitted with the Lsp1 crystal structure, albeit with unaccounted density at the lattice contact sites^29^. Our structures clearly reveal three regions of contact between the central dimer and its neighbours (Fig Ext Data 6A). The first site of contact is a short stretch of interactions between the well-folded, domain-swapped Nt of monomer A (res1-8) in the central dimer and the equivalent Nt stretch (res1-8) of monomer B in neighboring dimer 2, including residue S6 which previously shown to be phosphorylated by Pkh1 and important for eisosome assembly in combination with other phosphorylated residues^48,49^ (Fig Ext Data 6B). The remaining two contact sites are localized to the BAR domain tips, previously shown to be flexible in crystallographic studies^47^. A stretch of electrostatic interactions between residues 171-186 on helix 3, as well as residue 145 on helix 2, of the BAR domain in monomer A of the central dimer and the equivalent residues from monomer B of dimer 3 forms the second contact site (Fig Ext Data 6C). Finally, a hydrophobic interaction between Y155 at the tip of BAR domain helix 2 on monomer A of the central dimer with Y158 on monomer B of dimer 4, and vice versa, forms a third contact (Fig Ext Data 6D).

### Amphipathic helix associated with membrane void pattern

As mentioned above, immediately following the Nt lattice contact site is a previously unrecognized AH (res 39-48), oriented parallel to the BAR domain (Fig 1C). N-terminal AHs are a common feature of the N-BAR family of BAR domain proteins (e.g. endophilin, amphiphysin) and are proposed to play crucial roles in curvature sensing/induction. Insertion of an AH into the membrane expands only one leaflet, thus inducing local curvature via a “wedging mechanism”^35,50,51^. Additionally, amphipathic helices can recognize membrane curvature through their preferred insertion at sites of lipid packing defects induced by high local curvature^52^.

By making one-pixel slices through the unsharpened maps parallel to the axis of the eisosome tubule, we could clearly see the well-defined protein density of the AH buried within the cytoplasmic leaflet of the membrane (visible as a uniform density of lower intensity) (Fig 1D, panel 1, teal arrow). At slices ∼5Å deeper, an array of small voids in the membrane density begins to appear just below the AH. This pattern continues throughout the cytoplasmic leaflet, producing a fence-like striation in the membrane (Fig 1D, panels 2-5, violet arrows). However, this effect is asymmetric between the two leaflets: the pattern of voids is only present in the protein-bound leaflet, while the density of the exoplasmic leaflet is homogeneous throughout the lateral slices (Fig 1D, panels 6-10).

We wondered if these membrane voids could represent stably localized ergosterol molecules. Sterols, with their rigid ring structures, exhibit reduced Coulombic potential compared with tightly packed phospholipid tails^53–55^. Additionally, multiple examples of sterol/amphipathic helix interactions have been previously demonstrated. For example, transient interactions of sterols with the AH of Osh4 have been observed in MD simulations; cholesterol fills packing defects near aromatic side chains and wedges between acyl chains of poly-unsaturated PI(4)P^56^.

### Lipid binding pocket on Pil1/Lsp1 dimer

Visualization of the electrostatic surface of the membrane-facing surface of the Pil1 dimer revealed two patches of intense positive charge adjacent to the amphipathic helices (Fig 1E). Remarkably, within this pocket, an unassigned density is coordinated by several charged residues that have been previously proposed to be involved in PI(4,5)P_2_ binding^29,47^ (Fig 1F). *In vitro* tubulation by Pil1 and Lsp1 is reportedly dependent on the presence of PI(4,5)P_2_ in liposomes and defects in PI(4,5)P_2_ regulation cause changes in eisosome morphology *in vivo*^29,57,58^. Additionally, Lsp1 has been shown to cluster PI(4,5)P_2_ on giant unilamellar vesicles (GUVs) and prevent its lateral diffusion within the membrane^59^. Because of this well-documented structural and functional relationship between the eisosomes and PI(4,5)P_2_, we speculated that this unassigned density could represent one or more coordinated PI(4,5)P_2_ headgroups.

### Eisosome reconstitution with known lipids

In order to assign identities to the structural signatures we observe within the membrane, we chose to reconstitute eisosome filaments using lipid mixtures of known composition with recombinantly-expressed Pil1 protein. We tested several lipid mixtures with cholesterol and/or PI(4,5)P_2_, using as a constant component a mixture of 18:1 phosphatidylcholine (DOPC), 18:1 phosphatidylethanolamine (DOPE), and 18:1 phosphatidylserine (DOPS).

Our final reconstructions were made with three lipid mixtures: 1) “minus PI(4,5)P_2_/plus sterol”, containing 30% cholesterol 2) “plus PI(4,5)P_2_/minus sterol”, containing 10% PI(4,5)P_2_, and 3) “plus PI(4,5)P_2_/plus sterol”, containing 10% PI(4,5)P_2_ and 15% cholesterol (Supp Table 3&4). We used helical reconstruction to solve multiple structures of varying diameters, helical parameters, and resolutions. (Fig Ext Data 4B-D, 7A, Supp Table 1&2).

For all three lipid mixtures, the overall architecture of the Pil1 lattice is similar to the native samples. However, compared with the native samples, in “-PI(4,5)P_2_/+sterol” preparations, the amphipathic helices of the Pil1 dimers are not resolved (Fig 2A&B, Ext Data 3D, 7B). This could be because the AH is more mobile in this lipid mixture or because it is not inserted to the membrane. The AH (H_0_) of endophilin, for example, requires PI(4,5)P_2_ for membrane penetration^60^. In the “+ PI(4,5)P_2_/-sterol” samples the AH density is improved, while the resolution AH density is highest and most resembles that of the native eisosomes in the “+PI(4,5)P_2_/+sterol” samples, suggesting that PI(4,5)P_2_ is critical for AH stabilization (Fig 2C-D, Fig Ext Data 3E-F, 7C&D). To probe the PI(4,5)P_2_-dependency of the AH insertion into the membrane, coarse grained (CG) MD simulations were performed using a model tubule of Pil1 dimers and the exact lipid compositions of the reconstituted mixtures (“-PIP2/+sterol”, “+PIP2/-sterol”, and “+PIP2/+sterol”). The distances between the centre of mass (COM) of the AH and the head groups of the lipids were computed and, indeed, the averaged distance distributions show that in absence of PI(4,5)P_2_ the amphipathic helices partially detach from the membrane (Figure 2E, Fig Ext Data 8A-B).

**Figure 2.**
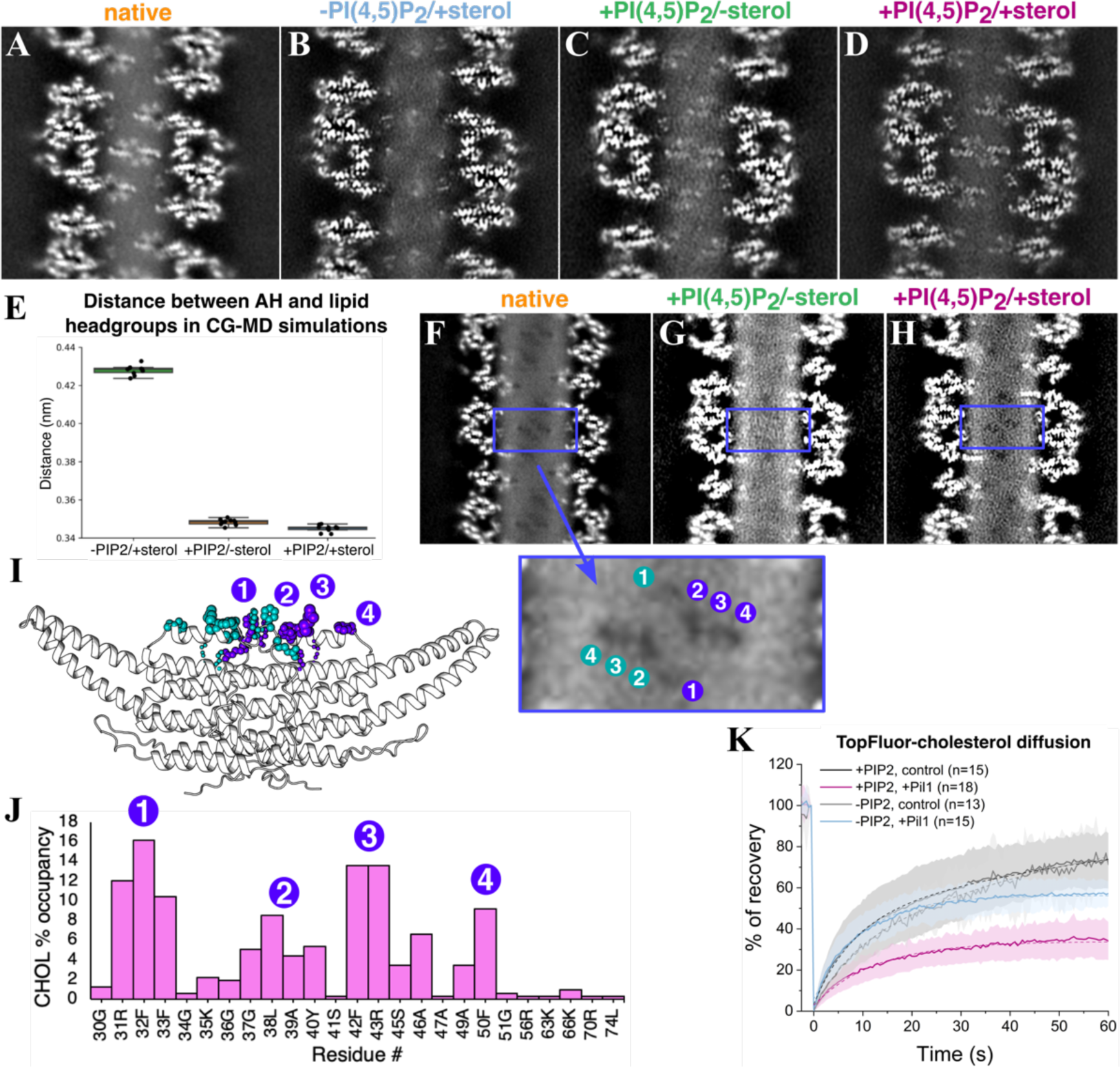
Sterols are stabilized by the Pil1/Lsp1 amphipathic helix within the eisosome membrane microdomain. **A-D.** Parallel slice at maximum amphipathic helix density of unsharpened maps of native (**A**), “-PI(4,5)P2/+sterol” reconstituted (**B**), “+PI(4,5)P2/-sterol” reconstituted (**C**), and “+PI(4,5)P2/+sterol” reconstituted (**D**) eisosomes. **E.** Distance between the center of mass of the AH and lipids’ head groups in CG simulations of different lipid compositions. **F-H.** Membrane voids pattern within the cytoplasmic leaflet in native (**F**), “+PI(4,5)P2/-sterol” reconstituted (**G**), and “+PI(4,5)P2/+sterol” reconstituted eisosomes (**H**). **I.** CG-MD snapshot showing AH-cholesterol interactions. Inset image: membrane voids with numbers indicating individual sterol dwell sites **J.** Occupancy of cholesterol at amphipathic helix residues in +PI(4,5)P2/+sterol system in CG-MD simulations. **K.** FRAP of TopFluor-cholesterol in control samples without protein and with Pil1 in presence and absence of 1% PI(4,5)P2. Solid lines indicate a mean of n number of measured nanotubes with standard deviation shown. Dashed lines indicate the fitted data.

### Membrane voids require PI(4,5)P_2_ and sterol

One feature that was notably absent from the “+PI(4,5)P_2_/-sterol” helices was the pattern of membrane voids in the cytoplasmic leaflet which we observe in the native eisosome filaments (Fig 2F-G). However, in the “+PI(4,5)P_2_/+sterol” helices, we can again observe the void pattern, supporting the notion that this pattern results from stable association of sterol molecules with the AH (Fig 2H). CG-MD simulations were performed with tubules of an identical lipid composition to the “+PI(4,5)P_2_/+sterol” mixture and the number of contacts between lipid headgroups and residues of Pil1 were measured for each lipid in terms of percentage of occupancy, i.e. the percentage of frames in which any lipid-protein contact is formed (Fig Ext Data 8C). Remarkably, sites of increased cholesterol occupancy were similar to the locations of the pattern of holes in the lipid density, with peaks at four clusters of residues, each containing aromatic sidechains: 1) residues 32F/33F, 2) residues 37G/38L/39A/40Y, 3) residues 42F/43R, and 4) residue 50F (Fig 2I-J). To better understand the relationship between PI(4,5)P_2_ binding and sterol dynamics, we reconstituted Pil1 scaffolds on pre-formed membrane lipid nanotubes to perform Fluorescence Recovery After Photobleaching (FRAP) assays using TopFluor-cholesterol in lipid mixtures with or without 1% PI(4,5)P_2_ (Supp Video 1). For nanotubes with Pil1 but without PI(4,5)P_2_ (-PI(4,5)P_2_/+sterol), there is a slight reduction in the mobile fraction of sterols, but this effect was pronounced in presence of Pil1 and 1% PI(4,5)P_2_ (+1% PI(4,5)P_2_/+sterol) (Fig 2K, Supp Table 5). This indicates that, in the presence of PI(4,5)P_2_, the immobilized fraction of cholesterol interacts strongly with either Pil1 and/or other lipids bound to the protein. Notably, the Pil1 lattice remained highly stable on the nanotubes without PI(4,5)P_2_ (Supp Video 2) indicating that the difference in sterol mobility is not due to unstable protein assembly.

### Lipid binding site in reconstituted eisosomes

In the “+PI(4,5)P_2_/-sterol” structures, a clear triangular density, which we fitted with an inositol-1,4,5-phosphate (IP_3_) ligand to represent a PI(4,5)P_2_ headgroup, was interacting with basic residues R126, K130, and R133 on each protomer in the binding pocket we previously identified in the native structures (Fig 3A, Ext Data 3E). Interestingly, these conserved residues were previously shown to be important for eisosome assembly *in vivo* and membrane binding *in vitro*^29,47^. Comparing this additional density with the putative lipid density observed in the native eisosome filament, there is a clear overlap (Fig 3A). This extra density, not present in the “-PI(4,5)P_2_/+sterol” Pil1 filaments (Fig Ext Data 3D), likely reflects a PI(4,5)P_2_ binding site in the native eisosomes. To confirm direct PI(4,5)P_2_ and Pil1 interaction, we again utilized pre-formed membrane lipid nanotubes, in this case to measure lipid sorting coefficients of 1% TopFluor-PI(4,5)P_2_ using a fluorescence ratiometric comparison with a reference lipid (Atto 647 N DOPE). This revealed a relative accumulation of PI(4,5)P_2_ in regions with Pil1 scaffolds (Fig 3B, Fig Ext Data 9A-D). We also checked TopFluor-PI(4,5)P_2_ diffusion in lipid nanotubes using FRAP assays and observed much slower recovery of PI(4,5)P_2_ when Pil1 is bound, suggesting a strong interaction (Fig 3C, Supp Table 5). A similar observation has been previously reported with Lsp1^59^.

**Figure 3.**
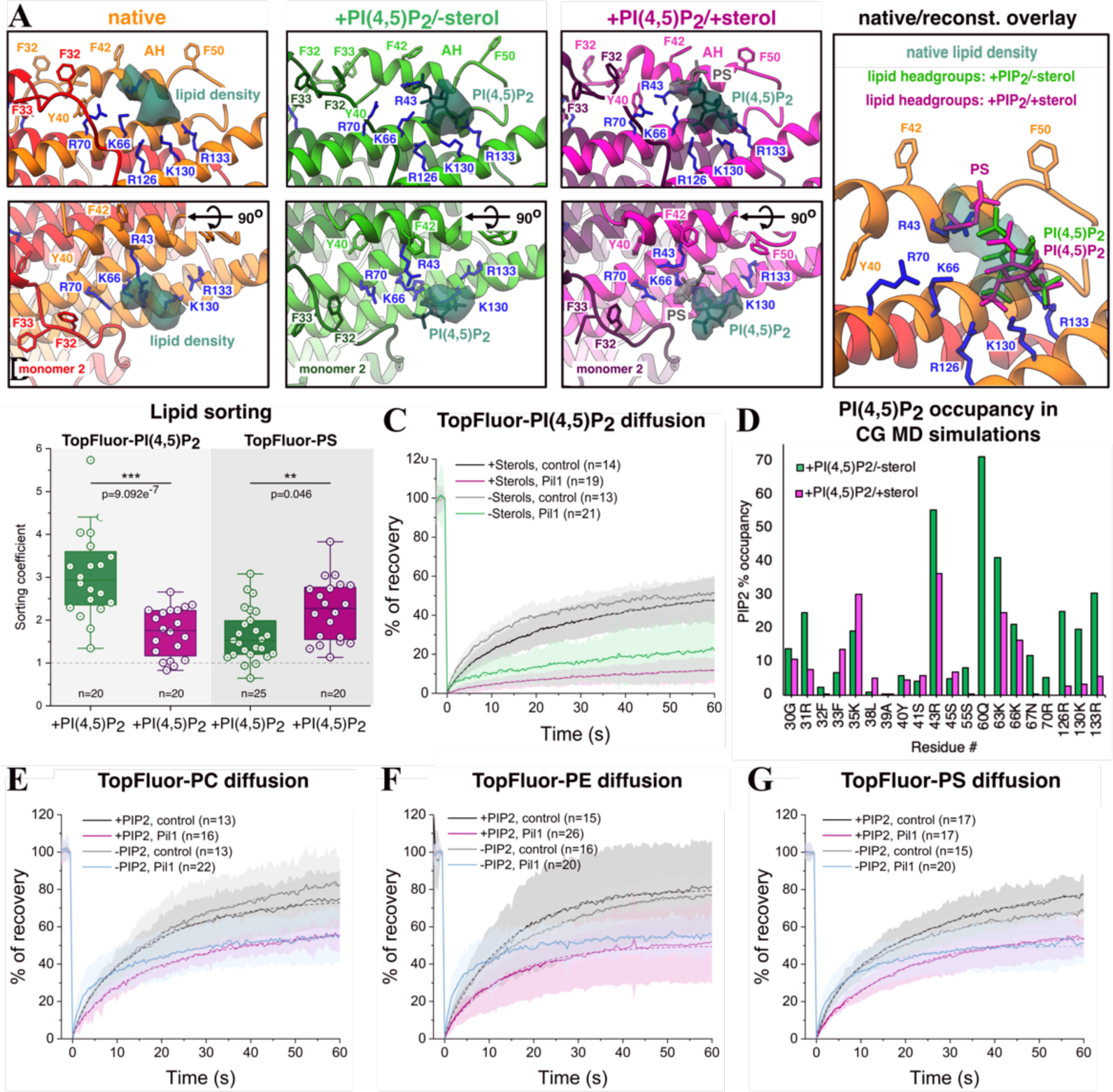
Reconstitution of purified Pil1 with lipids of known composition enables identification of structural signatures in native membrane. **A.** Lipid headgroup densities (sea green for PI(4,5)P2, grey for PS) in lipid binding pockets of native (orange), “+PI(4,5)P2/-sterol” reconstituted (lime green), and “+PI(4,5)P2/+sterol” reconstituted (magenta) eisosome maps. Rightmost panel: overlay of native eisosome density with fitted model of IP3 in “+PI(4,5)P2/-sterol” reconstituted structure (lipid headgroups indicated in lime green) and IP3 plus phospho-serine of “+PI(4,5)P2/+sterol” structure (lipid headgroups indicated in magenta). **B.** Lipid sorting co-efficients of PI(4,5)P2 and DOPS in “+1% PI(4,5)P2/-sterol” and “+1% PI(4,5)P2/+sterol” reconstituted Pil1 tubules. [Statistical significance: Tukeýs HSD test following one-way ANOVA assuming normal distribution, \*\*\**p* < 0.001, \*\**p* < 0.01, N=2]. **C**. FRAP of TopFluor-PI(4,5)P2 in “+1% PI(4,5)P2/-sterol” and “+1% PI(4,5)P2/+sterol” reconstituted Pil1 tubules. **D.** PI(4,5)P2 lipid occupancy for residues <5Å from PI(4,5)P2 headgroup with >5% occupancy in CG-MD simulations. **E-G.** FRAP of TopFluor-PC (**E**), TopFluor-PE (**F**) and TopFluor-PS (**G**) in control samples without protein and with Pil1 in “+1% PI(4,5)P2/-sterol” and “+1% PI(4,5)P2/+sterol” lipid nanotubes. Solid lines indicate a mean of n number of measured nanotubes with standard deviation shown. Dashed lines indicate the fitted data.

In the “+PI(4,5)P_2_/+sterol” samples, a similar triangular density bound to R126, K130, and R133 was observed and fitted with an IP_3_ ligand, comparable to the PI(4,5)P_2_ headgroup in “+PI(4,5)P_2_/-sterol” samples; however, parallel slices through the AH region reveal a slight decrease in intensity of this PI(4,5)P_2_ density in the “+PI(4,5)P_2_/+sterol” samples (Fig 3A and Ext Data 9E). In line with this observation, lipid sorting values for PI(4,5)P_2_ in the “+1% PI(4,5)P_2_/+sterol” samples revealed PI(4,5)P_2_ accumulation under Pil1 scaffolds, albeit to a lesser extent compared with “+1% PI(4,5)P_2_/-sterol” sample (Fig 3B). Additionally, in the CG-MD simulations for both the “+PI(4,5)P_2_/-sterol” and “+PI(4,5)P2/+sterol” systems, PI(4,5)P_2_ occupancy was increased at charged residues in the lipid binding pocket, especially residues R43, Q60, R126, K130 and R133, while addition of cholesterol reduced occupancy for all these residues, complementing the lipid sorting observations (Fig 3D, Fig Ext Data 8D). Curiously, using FRAP, we actually observed a reduction in PI(4,5)P_2_ mobility under the Pil1 scaffold in samples with cholesterol, suggesting that for the fraction of PI(4,5)P_2_ that is immobilized (e.g. protein-bound), sterols might play a role in enhancing the Pil1-PI(4,5)P_2_ interaction and perhaps stabilizing the AH (Fig 3C, Supp Table 5).

In the “+PI(4,5)P_2_/+sterol” samples, we were surprised to observe an additional small density stabilized between the AH and the PI(4,5)P_2_ headgroup, potentially coordinated by residues R43, K66, and/or R70 (Fig 3A). We speculated that this could be the signature of a lipid headgroup from one of the other lipids in our mixture. Phosphatidylserine (PS), in particular, is reported to strongly partition with sterols both *in vitro* and *in vivo*^61–63^. Using lipid nanotubes with TopFluor-labeled PS, we observed PS sorting in both the “+1% PI(4,5)P_2_/-sterol” lipid mixture and the “+1% PI(4,5)P2/+sterol” mixture, with a significantly higher sorting coefficient in the presence of sterol, supporting the notion that our extra density is a PS headgroup (Fig 3B). CG MD simulations also suggest that DOPS exhibits a similar affinity to the charged lipid-binding pocket of Pil1 as PI(4,5)P_2_, but with alteration (either enhancement or reduction) of several interactions with charged residues in the lipid binding pocket upon the inclusion of sterols in the lipid mixture (Fig Ext Data 8E-F). Consequently, we fitted this density with a phospho-serine ligand to represent a PS headgroup (Fig 3A). Collectively, these data suggest that the inclusion of sterol in the lipid mixture increases the specificity of PI(4,5)P_2_ and DOPS interaction with particular charged residues within the lipid binding pocket.

Remarkably, an overall reduction in lipid dynamics was also observed by FRAP for TopFluor-PC, -PE, and -PS with 1% PI(4,5)P_2_ and 30% cholesterol included in the membrane (Fig 3E-G). A significant portion of each of these lipids were immobilized in the presence of Pil1 indicating that the membrane is less diffusive under the Pil1 scaffold. While the fraction of these lipids that is immobilized by Pil1 is similar with or without PI(4,5)P_2_, the dynamics of the mobile lipid fraction for each of these lipids were decreased in the presence of PI(4,5)P_2_, again highlighting that PI(4,5)P_2_ is crucial for formation of stable lipid microdomains under the Pil1 scaffold (Fig 3E-G, Supp Table 5). This implies that the binding of PI(4,5)P_2_ and/or the stabilization of the AH of Pil1 create a sieve-like lattice within the cytoplasmic leaflet that slows lipid dynamics in the membrane microdomain, enabling the sterol molecules to form semi-stable interactions with bulky side chains of the AH.

### Physiological implications of Pil1 lipid binding

To understand the *in vivo* relevance of the lipid binding we observe, we produced mutant yeast strains that were predicted to affect the binding of Pil1 to different lipid species based on our structures and MD simulations (Fig 4A, Supp Table 6). Specifically, we tagged endogenous *PIL1* with *GFPenvy* in an *LSP1* deletion background and additionally introduced mutations to disrupt PI(4,5)P_2_ binding (*pil1^R^*^126^*^A^* and *pil1^K^*^130^*^A/R^*^133^*^A^*), PS binding (*pil1^R43A^* and *pil1^K66A/R70A^*), and both PI(4,5)P_2_ and PS binding (*pil1^K66A/R70A/K^*^130^*^A/R^*^133^*^A^*). We also produced a strain with mutations in the codons for the aromatic residues from each of the four sterol occupancy peaks observed in our CG-MD simulations (*pil1^F33A/Y40A/F42A/F50A^*) to disturb sterol localization.

**Figure 4.**
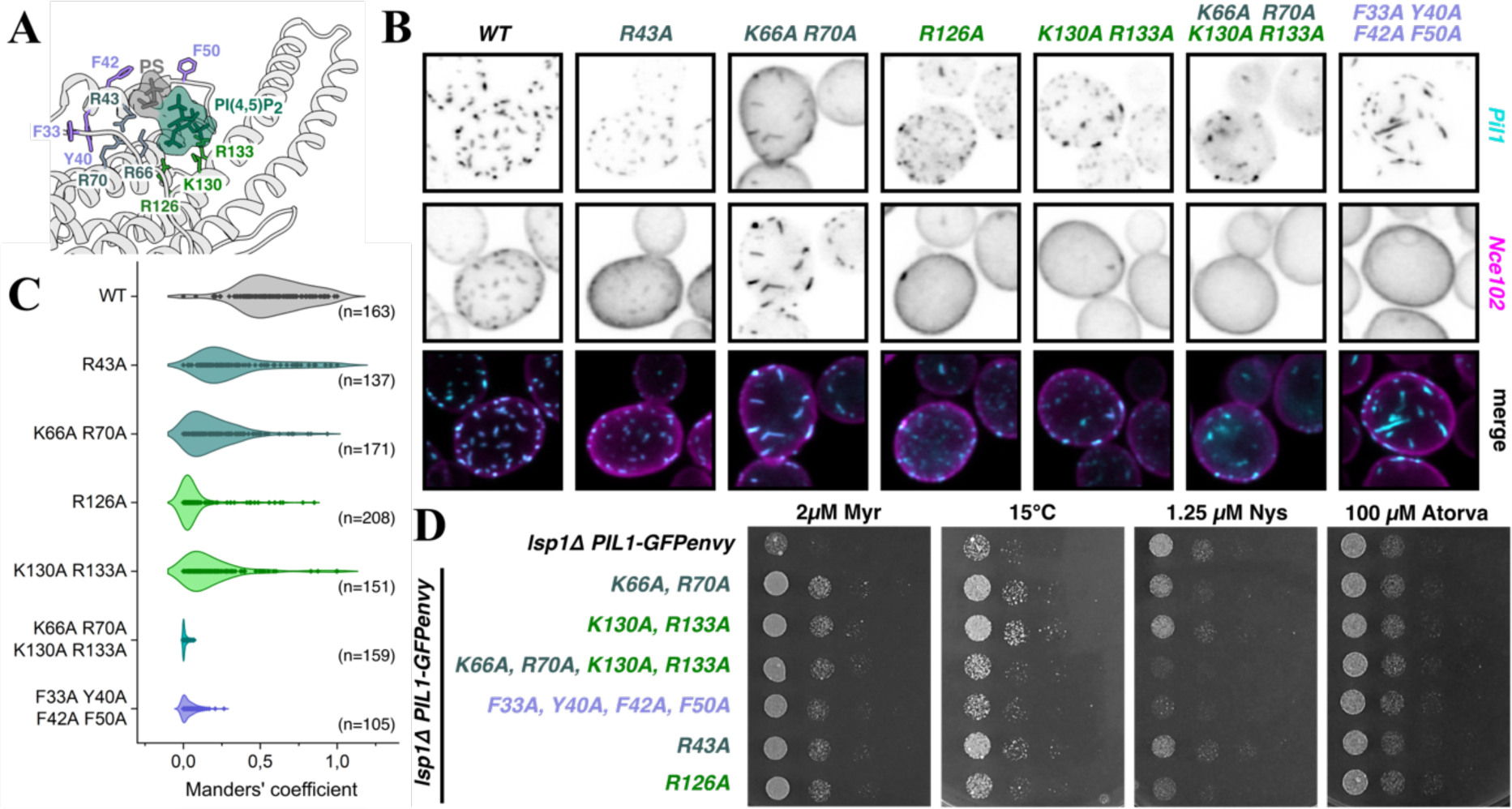
Lipid binding-impaired mutants affect morphology and function of eisosomes *in vivo*. **A.** A cartoon of the lipid binding pocket in the native sample. Sterol binding residues in violet, proposed PS binding residues in grey, PI(4,5)P2 binding residues in sea green. **B.** Eisosome morphology in *lsp1τι* yeast expressing, from their endogenous locus, Pil1-GFP with indicated lipid-binding mutations and Nce102-mScarlet-I (summed stacks). Merged represents summed stacks of Pil1-GFP (cyan) and mScarlet-I (magenta) signals **C.** Manders’ thresholded fraction of Nce102-mScarlet-I that colocalizes with Pil1-GFP lipid binding-impaired mutants in single cells (Manders’ M1 colocalization coefficient). The shaded area represents the probability for datapoints of the population to take on this value. **D.** Growth assays of *lsp1*τι yeast expressing Pil1-GFP lipid binding-impaired mutants. Myr: myriocin, Nys: nystatin, Atorv: atorvastatin.

Eisosome morphology and function was altered in all these mutants (Fig. 4B). Strains expressing *pil1^K66A/R70A^*, *pil1^K^*^130^*^A/R^*^133^*^A^*, and to a lesser degree *pil1^R43A^*, exhibited fewer eisosomes, but with an elongated morphology, while the *pil1^R^*^126^*^A^* and *pil1^K66A/R70A/K^*^130^*^A/R^*^133^*^A^* mutants displayed misshapen eisosomes and partial cytosolic mislocalization of the Pil1 protein. The *pil1^F33A/Y40A/F42A/F50A^* sterol binding-impaired mutants display unusual rod-like eisosomes that ingress towards the centre of the cell (Fig 4B, Fig Ext Data 9F). To further probe the role of these lipid binding sites, we chose to add an endogenous mScarlet-I tag to the eisosome resident protein Nce102. Nce102 re-localizes from the eisosomes into the bulk membrane (or vice versa) upon exposure to various membrane stressors including osmotic shocks, sphingolipid synthesis inhibition by the drug myriocin, and sphingoid base treatment^5,64,65^. Remarkably, all of the lipid binding-impaired mutants tested show a mislocalization of Nce102 already under steady state conditions, with nearly no co-localization in the *pil1^K66A/R70A/K^*^130^*^A/R^*^133^*^A^* (PI(4,5)P_2_ and PS binding-impaired) and *pil1^F33A/Y40A/F42A/F50A^* (sterol binding-impaired) mutants (Fig 4C).

To determine if there are physiological consequences triggered by loss of lipid binding, we assessed growth of our mutant strains under a variety of stress conditions. All of our Pil1 lipid binding-impaired mutants exhibited myriocin-resistance (Fig 4D, Fig Ext Data 9G). This is in line with the previously described role of eisosomes in sphingolipid homeostasis^64,66^, and similar to what has been previously reported for a mutant lacking Pil1 and all eisosomal tetraspan proteins of the Nce102 and Sur7 families^67^. Interestingly, we observed that *pil1^K^*^130^*^A/R^*^133^*^A^*, and to a lesser degree *pil1^R43A^* and *pil1^K66A/R70A^*, grew better at low temperature relative to control. As cold resistance has been previously described for cells lacking the eisosome-localized PI(4,5)P_2_ phosphatase Inp51, or Inp52^68^, it is possible that this cold resistance could be caused by either PI(4,5)P_2_ dysregulation or Inp51/52 mislocalization.

We also checked growth with nystatin, an antimycotic that interacts with plasma membrane free ergosterol to induce cell lysis^69,70^. Strikingly, the sterol binding-impaired *pil1^F33A/Y40A/F42A/F50A^* mutant and the *pil1^R^*^126^*^A^* and *pil1^K66A/R70A/K^*^130^*^A/R^*^133^*^A^* mutants which exhibited cytosolic mislocalization were sensitive to nystatin, indicating a higher availability of free ergosterol at the plasma membrane (Fig 4D, Fig Ext Data 9G). However, none of the mutants showed resistance to the sterol synthesis inhibitor atorvastatin, indicating the nystatin sensitivity is not due to enhanced ergosterol synthesis^71^. It is tempting to speculate that that these mutations affect the lipid microdomain generated by Pil1, rendering a normally sequestered population of ergosterol available for nystatin binding, though it is also possible that a more general disorganization of ergosterol localization occurs due to eisosome-dependent signaling dysregulation.

### 3D variability analysis of native eisosome lattice

With a clear understanding of the identities of the lipids that produce the signatures we see within the membrane and their functional importance, we returned to the native eisosome dataset to check for variability in the lipid organization. Using the symmetry expanded/density subtracted lattice particles, we performed 3D variability analysis. One component of our 3D variability analysis was particularly striking; we observed conformational flexibility within the Pil1 lattice that resembled a spring-like stretching and compression at the Nt contact sites (Supp Video 3). Because the AH is directly connected to this Nt region, we considered whether this stretching could be transmitted to the lipid bilayer. Remarkably, we could see that changes in the shape and size of the bound lipid density were synchronized with the stretching of the Nt contact sites.

To understand the relationship between the Nt stretching and the changes in the lipid density, we extracted 10 sets of non-overlapping particles along this variability dimension for further refinement (Fig Ext Data 4E-F, 10A). For these 10 classes, we also analyzed the tubule of origin for each particle to determine the contribution of tubule diameter to this stretching component. Particles from every tubule diameter are distributed across all 10 classes; however, particles from small diameter tubules are over-represented in the more compact classes, while those from large diameter tubules are over-represented in more stretched classes (Fig Ext Data 10B-C).

In the deepEMhancer sharpened maps, the putative lipid binding pocket of the most compressed class contained two clear densities, one triangular density that we could identify as a PI(4,5)P_2_ headgroup with interactions at residues R126, K130, and R133, and the other smaller elongated density interacting with residue K66, which we had previously assigned as a PS headgroup in the “+PI(4,5)P_2_/+sterol” reconstituted samples (Fig 5A, Fig Ext Data 3G). In contrast, the lipid binding pocket in the most stretched class was unoccupied in the sharpened maps, suggesting PI(4,5)P_2_ and PS binding are disrupted upon stretching (Fig 5B, Fig Ext Data 3H).

**Figure 5.**
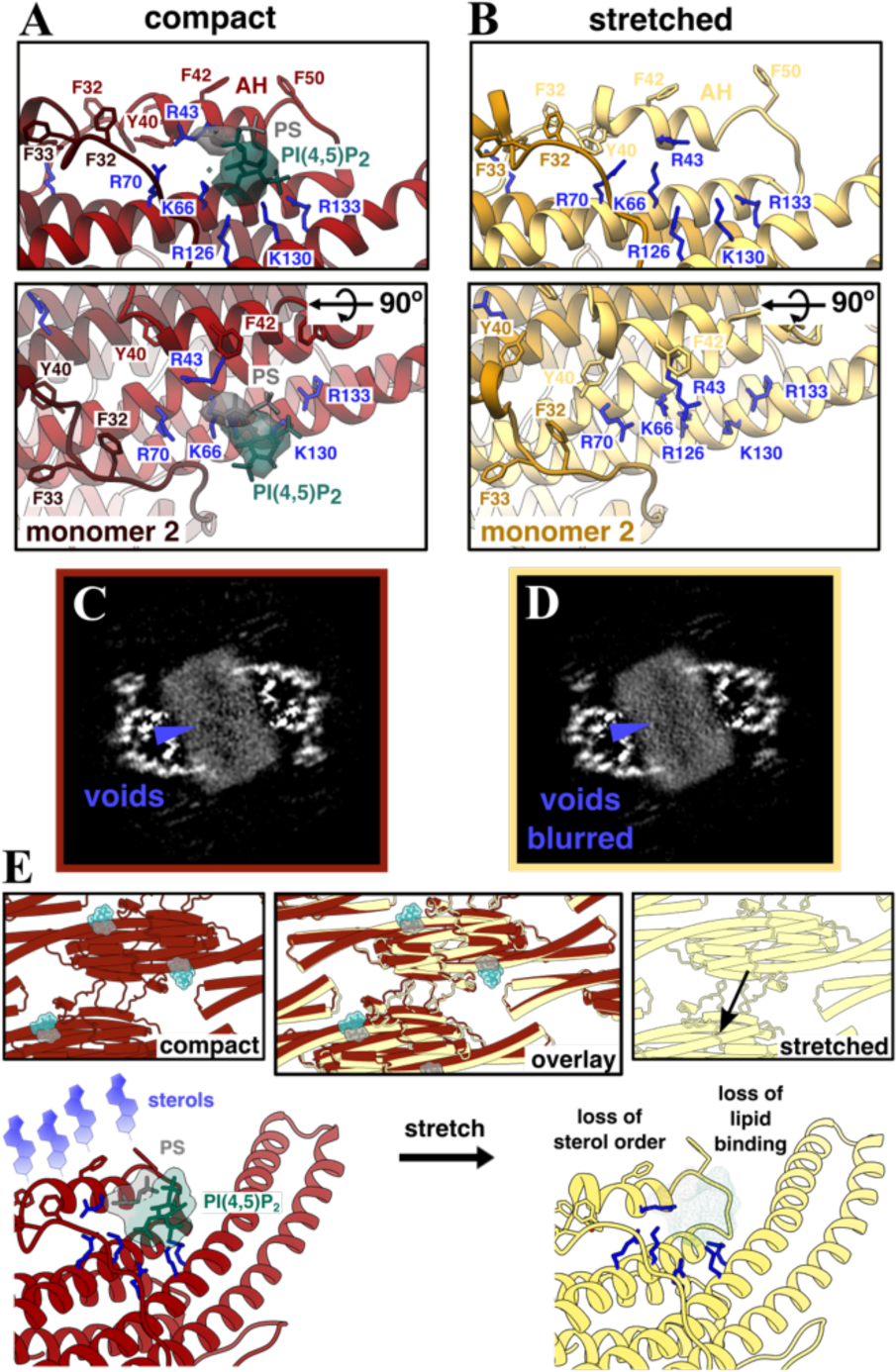
3D variability analysis reveals Pil1/Lsp1 lattice stretching and its effects on membrane organization within native eisosomes. **A**. Well-defined two-part density (sea green for PI(4,5)P2, grey for PS) occupying charged pocket in deepEMhancer sharpened map of most compact lattice conformation (model in dark red). **B.** No clear density observed in charged pocket in sharpened map of most stretched conformation (model in light yellow). **C-D.** Membrane void pattern, or lack thereof, in corresponding slices of unsharpened maps of most compact (**C**) and most stretched (**D**) conformation. **E.** Model illustrating membrane stretching destabilizes lipid headgroup and sterol binding (compact model in dark red, stretched model in light yellow, PI(4,5)P2 headgroup in sea green, PS headgroup in grey, sterols in violet).

To further investigate the differences between these classes, we analyzed slices through the membrane of the unsharpened maps. Strikingly, while the most compressed class retained the pattern of membrane voids beneath the AH, indicating sterol binding, this pattern was blurred in the most stretched class, despite its slightly higher resolution (Fig 5C-D, Ext Data 10D-G, Supp Table 1). However, no overall changes in membrane thickness or overall intensity were observed in radial angle profile plots of these two classes (Fig Ext Data 10H). This suggests that sterols are mobilized rather than redistributed in the stretched class.

## Discussion

Taken together, our observations provide clear evidence that the eisosome scaffold proteins Pil1/Lsp1 form a plasma membrane microdomain via their direct interactions with specific lipids. We believe this work provides an unprecedented level of detail into the organization and dynamics of plasma membrane lipids within this microdomain. Furthermore, our data allow us to propose a speculative model for how these eisosome microdomains could sense and respond to mechanical stress.

While the Pil1/Lsp1 lattice is capable of self-organization on the membrane surface in the absence of PI(4,5)P_2_, we find that PI(4,5)P_2_ is necessary for the stabilization of the AH within the cytosolic leaflet. When PI(4,5)P_2_ is bound and the AH is inserted, semi-stable interactions of sterols, mostly with bulky side-chains of the AH, are able to form and the mobility of lipids (and likely the eisosome-resident proteins, like Nce102) within the membrane microdomain is reduced. Notably, the stabilized lipid interactions we observe are limited to the cytoplasmic leaflet, demonstrating clear membrane asymmetry in the intra-leaflet lipid dynamics.

Under mechanical stress, Pil1/Lsp1 lattice stretching could conceivably be communicated to the lipids in the cytoplasmic leaflet through the direct connection between the Nt lattice contact sites and the AH. In our native structures, we see that lattice stretching is correlated with a mobilization of PI(4,5)P_2_ and a loss of the sterol patterning within the cytoplasmic leaflet (Fig 5E), which implies an increased mobility of all of the lipids within the eisosome. Given that all of our Pil1 lipid binding-impaired mutants, even those with mild morphology defects, exhibit mislocalization of the tension-responsive protein Nce102, we propose that this lipid mobilization represents a general mechanism to free sequestered factors to initiate their membrane stress-sensitive signaling functions. These sequestered factors need not be proteins; for example, our mutants predicted to be compromised in sterol coordination are hypersensitive to Nystatin, a macrolide antibiotic that binds sterols in the plasma membrane leading to cell leakage and death, suggesting that sequestration of sterol at eisosomes can also regulate the amount of free sterol in the PM. Lastly, although we have presented eisosome lattice stretching as a possible trigger leading to release of sequestered factors, other membrane perturbations including changes in lipid dynamics triggered, for example, by thermal shock or amphiphilic toxins, could also be sensed by this system.

This remarkable function of AHs to organize the lipids within the membrane and connect mechanical stretching to the dynamics of those lipids may be a conserved feature of mechanosensitive proteins. In fact, AHs from many mechanosensitive ion channels, including Piezo1^72–74^, Piezo2^75^, TREK/TRAAK^76,77^ and OSCA1.2^78^ have bulky side chains inserted to the membrane and Piezos, in particular, have been proposed to modulate and be sensitive to the lipid composition of the membranes in which they function^79–81^.

Although the eisosomes are a fungi-specific membrane feature, many of the principles of lipid coordination by BAR domain proteins and other proteins that form lattices on membrane surfaces are likely to be conserved. For example, caveolae in mammalian cells, scaffolded by the interaction between caveolins (integral membrane proteins) and cavins (peripheral membrane proteins) are proposed to concentrate cholesterol, PS, and PI(4,5)P_2_^31,82,83^, as well as flatten in response to sterol removal from the plasma membrane^84^, suggesting that they share many features with the eisosome membrane microdomain. In fact, the organization and alteration of the dynamics of membrane lipids through interactions between membrane scaffolding proteins, charged lipids, and sterols could extend across a wide variety of structurally and functionally diverse proteins that have been proposed to associate with membrane microdomains, from focal adhesion proteins^85,86^ to ESCRTs^87,88^ to myelin-specific proteins^89,90^. Our detailed characterization of the eisosome membrane microdomain provides a novel context for understanding how protein-lipid interactions participate in cell signaling functions.

## Methods

### Yeast strains

Yeast strains used for endogenous protein purification were constructed using classical recombination methods. Yeast strains with point mutations were constructed using CRISPR-Cas9 based methods. Strains used in this study are listed in Supplementary Table 6.

### Native eisosome purification

Yeast expressing Bit61-TAP from the endogenous locus were grown to optical density (OD_600_) of 6-8, harvested by centrifugation at 6000rpm for 10 mins, flash frozen in liquid nitrogen, and stored at -80°C. Cells were lysed by manual grinding with mortar and pestle under liquid nitrogen, then resuspended by slow rotation at 4°C in 1.5 volumes of extraction buffer (50mM PIPES pH 7, 300mM NaCl, 0.5mM CHAPS, 0.5mM DTT plus 1mM PMSF and 1× Complete protease inhibitor cocktail (-EDTA) (Roche)). Lysates were cleared by centrifugation at 12000rpm (xg) for 10 mins, and supernatants were incubated with IgG-coupled Dynabeads M270 (ThermoFisher Scientific) for 2 hrs at 4°C. Beads were washed 5 times with wash buffer (50mM PIPES pH 7, 300mM NaCl, 1mM CHAPS, 0.5mM DTT) at 4°C, then incubated with TEV protease (0.1 mg/ml) for 1hr at 18°C. Eluate was collected at 4°C then used immediately for cryoEM grids preparation.

### Plasmids and protein purification

*PIL1* was cloned into pCoofy6 vector (a gift from Sabine Suppmann, Addgene plasmid # 43990) as described in Scholz et al.^91^ (doi: 10.1186/1472-6750-13-12) with following primers: LP1-Sumo3-Pil1-fwd 5’-gtgttccagcagcagaccggtggaatgcacagaacttactctttaag, LP2-ccdB-Pil1-rev 5’-ccccagaacatcaggttaatggcgttaagctgttgtttgttggggaag, LP2-ccdB-fwd 5’-cgccattaacctgatgttctgggg, and LP1-Sumo3-rev 5’-tccaccggtctgctgctggaacac. After PCR-amplification using Q5 High-Fidelity 2X Mastermix (#M0492, New England Biolabs Inc.) gel-purified DNA fragments were assembled using RecA recombinase (#M0249, New England Biolabs Inc.). A plasmid containing mCherry was a kind gift from Alphée Michelot (Aix-Marseille Université). The mCherry was cloned into the C-terminus of Pil1 with following primers: LP1-Pil1-linker-mCherry-fwd 5’-tctcttccccaacaaacaacagctgagctcgctgcagcaatggtgag, LP2-mCherry-rev 5’-tggtgctcgagtgcggccgcaagcctagtttccggacttgtacagctc, LP1-Pil1-rev 5’-agctgttgtttgttggggaagagac, and LP2-pCoofy-vector-fwd 5’-gcttgcggccgcactcgagcaccac. The DNA fragments were amplified and assembled as above.

Recombinant Pil1 and Pil1-mCherry were expressed in BL21(DE3)pLysS (#200132, Agilent Technologies Inc.) in auto induction LB medium (#AIMLB0210, Formedium Ltd.) overnight at 20°C. Cells were lysed in lysis buffer (20 mM HEPES, pH 7.4, 150 mM KCl, 2 mM MgAc, 30 mM imidazole) supplemented with 1% Triton X-100, 1 mM PMSF and cOmplete protease inhibitor cocktail (#5056489001, Roche) by sonication on ice. Proteins were first purified with HisTrap Fast Flow column (#GE17-5255-01, Cytiva) in Äkta Pure system (Cytiva) using gradient of imidazole from 30 mM to 500 mM. Proteins were subsequently dialyzed overnight with 20 mM HEPES, pH 7.4, 75 mM KCl, 2 mM MgAc buffer and further purified with HiTrap Q sepharose HP column (#17115401, Cytiva). A KCl gradient from 75 mM to 500 mM was used to elute the protein. To cleave off the Sumo3 tag, SenP2 protease in final concentration of 30 µg/ml was then added and protein were dialyzed with 20 mM HEPES, pH 7.4, 150 mM KoAc, 2 mM MgAc buffer overnight. Finally, protein samples were cleaned with Superdex 200 Increase 10/300 GL column (#GE28-9909-44, Cytiva) equilibrated with 20 mM HEPES, pH 7.4, 150 mM KoAc, 2 mM MgAc buffer. Proteins were concentrated to 20-25 µM, snap-frozen with liquid nitrogen and stored in -80°C.

### Reconstitution of Pil1 tubules

Lipids used are listed in Supplemental Table 3. To reconstitute Pil1 tubules using large unilamellar vesicles (LUVs) for cryoEM, lipids were mixed in chloroform to final concentration of 3.8 mM with desired molar ratios. Chloroform was evaporated under argon gas flow and subsequently for three hours in 30°C vacuum oven. A lipid film was hydrated in reaction buffer (20 mM HEPES, pH 7.4, 150 mM KoAc, 2 mM MgAc), subjected to 10 cycles of freeze-thaw and extruded through a 200 nm pore-sized polycarbonate filter (Cytiva Inc.) using a mini-extruder (Avanti Polar Lipids Inc.). Lipid compositions used in cryoEM studies are listed in Supplemental Table 4. To produce samples for cryoEM studies, a mixture of 15-20uM recombinant Pil1 and ∼2mg/mL LUVs were incubated at 30°C for 1hr before freezing.

To reconstitute Pil1 tubules on pre-formed membrane nanotubes for fluorescence microscopy experiments, supported lipid films over silica beads were formed from multilamellar vesicles by mixing lipids in chloroform to final concentration of 1 mg/ml and evaporating the solvent as described previously^92^. Briefly, lipid films were hydrated using 5 mM HEPES, pH 7.4 buffer. Multilamellar vesicles were then mixed with silicon dioxide microspheres (Corpuscular #140256-10 or Sigma-Aldrich #904384) and dried for 30 minutes in 30°C vacuum oven. Imaging chamber was prepared by attaching a sticky-Slide VI 0.4 (Ibidi GmbH, #80608) on a 24 × 60 mm microscope cover glass. Sample chambers were passivated for 10 minutes with 2 g/l bovine serum albumin solution and subsequently washed several times with reaction buffer (20 mM HEPES, pH 7.4, 150 mM KoAc, 2 mM MgAc). Lipid-coated silica beads were then hydrated by adding a small amount of beads in the sample chamber and allowing beads to roll through the chamber. Several different lipid compositions were used in these experiments (See Supplemental Table 4). 0.01 mol% of Atto647N DOPE was added to each lipid mixture to visualize nanotubes and as the reference fluorescent lipid for lipid sorting coefficient measurements.

### CryoEM grid preparation and data collection

5ul of fresh sample was applied to untreated Lacey Carbon film on copper mesh grids (Jena Bioscience #X-170-CU400), blotted for 3-4s, then re-applied, blotted for 2-4s (2nd blot), and finally plunge-frozen in a Leica GP2 plunge system at 18oC, 90% humidity.

Native eisosome filaments were imaged by targeted acquisition using SerialEM with a 300kV Titan Krios fitted with a Gatan K2 Quantum direct electron detector (Heidelberg). A total of 2827 movies were collected, each with a total dose of 40 e-/Å^2^, a target defocus range of -0.8 to -1.8 μm and a pixel size of 1.327Å (105kx magnification).

Reconstituted Pil1 filaments were imaged using EPU software with a 300kV Titan Krios and a Falcon 4 direct detector (DCI Lausanne). Three datasets were collected: 1) Pil1 + “minus PI(4,5)P2” liposomes (21,386 movies), 2. Pil1 + “minus cholesterol” liposomes (22,960 movies), and 3) Pil1 + “PI(4,5)P2/cholesterol” liposomes (22,408 movies). For each movie, a total dose of 50 e-/Å2, a target defocus range of -0.6 to -1.8 um and a pixel size of 0.83Å (96,000x magnification) was used.

### CryoEM data processing

The pipeline for cryoEM data processing is outlined in Extended Data Figures 1,2, and 7.

For native eisosomes, movies were aligned in using MotionCor2^93^ and CTF correction was completed using Gctf^94^. Filaments were handpicked using manual picking in RELION v2.1.0. 2D classification was run iteratively in RELION 2.1. For particles in each clean RELION 2D class, power spectra were summed by class, then manually sorted into identical “types”. Helixplorer-1^95^ was used to estimate helical parameters, which were used with particles from each “type” for 3D auto-refinement with helical parameters in RELION. All helix types were then corrected for handedness and aligned along the D symmetry axis (using C symmetry worsened resolution). A mask was generated in RELION covering the central third of the helix and a final round of helical refinement was completed either with the mask (to optimize resolution) or unmasked (to be used for particle subtraction). Resolution estimates for masked maps are based on gold standard FSC values with a 0.143 cutoff on post-processed maps with the one-third mask used for refinement, a manually chosen initial threshold, and auto-bfactor calculation.

Symmetry expansion and density subtraction on native filaments was completed with unmasked maps from the final iteration of 3D auto-refinement. Helical parameters for each helix type were used for symmetry expansion but using C symmetry instead of D symmetry. A mask for density subtraction was generated in Chimera v1.5 by the addition of two zone maps: 1) an 8Å zone using models of a central Pil1/Lsp1 dimer and the six dimers with which it shares lattice contact sites and 2) a spherical zone of 60Å centered on the amphipathic helices of the Pil1/Lsp1 dimer. This initial mask was then extended with a soft edge, then used for density subtraction and reboxing of the particles in RELION v3.1.3. Subtracted particles were then used to reconstruct a volume and particles for all helix “types” and the reconstructed volume were imported into cryoSPARC v4.1.2 for further processing. Homogenous refinement was completed with all particles with the reconstructed volume as the initial model using C2 symmetry. This map was used for refinement of the Pil1 and Lsp1 native models (See “Model building’ for details). This map was then symmetry expanded in C2 for 3D variability analysis. 3D variability analysis was completed with 5 components requested. Manual inspection revealed one component that exhibited lattice stretching. This component was used for 3D variability display in intermediate mode with 10 non-overlapping frames used to generate particle subsets. Each particle subset was then used for masked local refinement with a mask covering the central dimer (generated in Chimera using the Pil1/Lsp1 dimer with a 10Å zone and extended with a soft edge in RELION). These maps were used for refinement of the Pil1 compact and stretched (native) models (See “Model building’ for details). Sharpening with deepEMhancer^96^ was used to improve the resolution of lipid headgroups in the lipid binding pocket. Resolution estimates are based on gold standard FSC values with a 0.143 cutoff using an optimized mask automatically generated during refinement.

For reconstituted Pil1 filaments, all data processing was completed in cryoSPARC v4.1.2. Movies were processed on-the-fly with CryoSPARC Live v3.2.2, using patch motion correction and patch CTF estimation. Filaments were picked using Filament Tracer, then cleaned and sorted using iterative rounds of 2D classification. Clean classes were used to calculate average power spectra, which were then manually sorted into identical “types”. Helixplorer-1 was used to estimate helical parameters, which were used with particles from each “type” for helical refinement. After initial refinement all helix types were corrected for handedness and aligned along the D symmetry axis. A mask on the central third of the helix was created in Relion v3.1.3 and used for a final round of helical refinement. These maps were used for refinement of the Pil1 lattice (-PIP2/+sterol reconstituted), Pil1 lattice (+PIP2/-sterol reconstituted), and Pil1 lattice (+PIP2/+sterol reconstituted) models (See “Model building’ for details). Resolution estimates are based on gold standard FSC values with a 0.143 cutoff using an optimized mask automatically generated during refinement.

Figures were made in Chimera v1.16 or Chimera-X v1.5. Parallel slice images were made in FIJI using the Reslice tool without interpolation. 3D intensity plots were made in FIJI using the 3D surface plot tool on the slice with maximum sterol void intensity using identical display conditions on both maps. Radial angle profile plots were made in FIJI using the Radial Profile Extended plugin with an angle of 40 degrees, corresponding to the size of the sphere used in density subtraction to include the lipid density under the central dimer.

### Model building

Structure predictions for Pil1 and Lsp1 from the AlphaFold database (https://alphafold.ebi.ac.uk/) were used as starting models, with the C-terminal region removed, starting from residue 275, for which no density was observed. Iterative rounds of model building, performed in Coot v.0.8.9.2, and real-space refinement, performed with Phenix-1.20-4459, were completed until no improvement in the model was observed. The model quality and fit to density were performed using Molprobity and Phenix-1.20-4459.

Ligand constraints for inositol 2,4,5-triphosphate and the phospho-serine were produced using phenix.elbow. Refinements with ligands were performed with these ligand constraints. For lipid headgroup ligands in native eisosome compact dimer model, ligands refined in the Pil1 “+PI(4,5)P2/+sterol” reconstituted map were placed in the in the deepEMhancer sharpened native eisosome compact map, then adjusted in Coot with rigid body fitting.

An electrostatic potential map of the model surface was calculated using the Coulombic potential function of ChimeraX v1.5.

### Lipid diffusion measurements with Fluorescence Recovery After Photobleaching (FRAP)

Lipid nanotubes were prepared as described above and 200 nM of Pil1-mCherry was incubated with nanotubes for 30 minutes. FRAP experiments were performed with Olympus IX83 wide-field microscope equipped with an Olympus Uapo N 100x 1.49 oil objective and an ImageEM X2 EM-CCD camera (Hamamatsu). The system was controlled by the Visiview software (Visitron Systems GmbH). Five frames were captured before a small protein-coated region was bleached. Subsequently, the recovery of fluorescence intensity was measured by capturing images every 500 ms for 1-2 minutes. The FRAP data was analyzed with Fiji/ImageJ by measuring intensity over time from the bleached region (region-of-interest), the region on nanotube outside the photobleached region (bleaching correction) and the region outside nanotube (background). After background subtraction and bleaching correction, the data for normalized to the intensity values before photobleaching. Graphs were generated with Origin Pro 2022 (OriginLab Corp.). To calculate recovery halftimes and mobile fractions, we used either one-phase exponential equation:

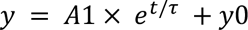

or two-phase exponential equation:

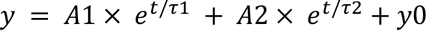

where A is value of y in plateau, t is time, τ is a time constant and y0 is the value of y when t=0.

Recovery halftimes were then calculated with equation:

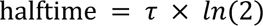

### Lipid sorting coefficient measurements

Lipid nanotubes were prepared as described earlier and incubated with 200-400 nM of Pil1-mCherry for 30 minutes until protein scaffolds were formed and visible by fluorescence microscopy. Imaging was performed using an inverted spinning disk microscope assembled by 3i (Intelligent Imaging Innovation) consisting of a Nikon Eclipse C1 base, a 100x 1.3 NA oil immersion objective. Sorting coefficients were calculated for each of the mentioned lipids using Atto 647N DOPE as the reference lipid with the following equation:

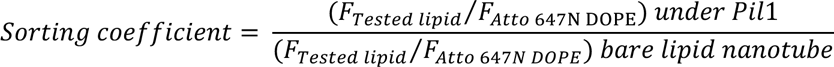

where *F_Tested lipid_* and *F_Atto 647N DOPE_* are the integrated fluorescence densities of the lipid problem and lipid reference integrated from the lipid nanotube plot profiles (See Fig Ext Data 9D) after background subtraction and neglecting the polarization factor^97^. Data and graphs were generated using Origin Pro 2022 (OriginLab Corp.).

### Molecular dynamics simulations

In accordance with the experimental models, three systems with different lipid compositions were modelled and simulated. Coarse grained (CG) molecular dynamics (MD) simulations of Pil1 tubule interacting with the tubule membranes were performed, in duplicates.

The tubule membranes were built using the BUMPy Software^98^. The systems, without the protein, were solvated with water and minimized using the steepest descent algorithm^99^. Four equilibration steps were performed, as follows: i) A first equilibration with a timestep of 5fs was run for 10ns, imposing position restraints with a force constant (fc) of 300kj/mol/nm2 on the lipid tails, to allow formation of membrane pores to equilibrate lipid and water content between the tubule lumen and the external region. 2) A second equilibration step of 5ns was performed using the previous settings but increasing the fc at 500kj/mol/nm2 and the timestep to 10fs. 3) A third equilibration step was run for additional 10ns after removal of position restraints to allow the closure of the pores. The Berendsen barostat and the v-rescale thermostat^100,101^ -with a temperature of 303K - were used.

From the CryoEM structure of Pil1 tubule (21 dimers), CG mapping was performed using Martinize2^102^, imposing an elastic network within the dimers. The CG protein model was then manually positioned around the tubule membrane using VMD v1.9^103^. The final system was solvated with water beads and neutralized adding Na+ and CL-ions. Each system was minimized and equilibrated in seven steps: 1) A first equilibration with a timestep of 5fs was run for 10ns, imposing position restraints with a force constant (fc) of 300kj/mol/nm2 on the lipid tails, to allow waterpore formation. The Berendsen barostat^100^ was applied to all the direction, with τp=5, and the v-rescale thermostat was used, setting the temperature at 303K^101^. 2) A second equilibration step of 5ns was performed using the previous setting but increasing the fc at 500kj/mol/nm2, and the timestep at 10fs. 3) A third equilibration step was run for other 10ns increasing the fc at 1000kj/mol/nm2, to maintain the waterpores open and to allow for the solvent equilibration. 4) Starting from the 4th step, the fc on the tails was progressively reduced to slowly induce a slowly closure of the pores. Finally, a fc of 500kj/mol/nm2 was applied, to run an equilibration of 5ns. 5) An equilibration decreasing the fc at 300kj/mol/nm2 was carried out for other 5 ns. 6) An equilibration removing the fc on the lipid tails was performed to allow the complete closure of waterpores. 7) A final equilibration step of 10ns was run without restraints, increasing the timesteps to 20fs.

For systems containing cholesterol the equilibration procedure was extended including an additional first equilibration step, with a reduced timestep of 2fs.

For production, the Parrinello-Rahman barostat^104^ was used with τ_p_=12. For each system, two replicates of approximately 10μs were performed. The simulations were carried out using the GROMACS Software v2021.5^99^ and the Martini3 force field^105,106^.

To compute the lipids’ occupancy, the PyLipID python package was used^107^. The analysis was performed selecting the head groups of lipids. The values were averaged over time and over the dimers. To investigate differences in the insertion of the amphipathic helices based on the different lipid membrane composition, the distances between the centre of mass (COM) of the amphipathic helices and the head groups of the lipids were calculated using gmx mindist tool of GROMACS 2021.5 Software^99^.

### Spot assays

Saturated overnight yeast cultures (30°C, SC medium) were diluted to OD_600_ 0.1 in the morning and grown into mid log phase (OD_600_ 0.5-0.8). Log phase cells were diluted to OD_600_ 0.1, and a 10-fold dilution series was spotted onto pre-dried SC media plates containing treatment substances, or vehicle. Plates were incubated at 30°C, except low (15°C) temperature plates, and imaged when differences were most apparent (typically after 24h for Nystatin, 48h for controls and Atorvastatin, 144h for myriocin, and 168h for 15°C). Substance stocks used in this study: Myriocin (Sigma M1177) 2.5mM in MeOH, Nystatin (Sigma 475914) 50mM in DMSO, Atorvastatin (Sigma PHR1422) 20mM in DMSO.

### Fluorescence microscopy and image evaluation

Logarithmically growing overnight yeast cultures (30°C, SC medium) were diluted and grown to OD_600_ 0.6. For Fluorescence live cell microscopy, cells were loaded into a Concanavalin coated flow chamber (ibidi μ-Slides VI 0.4 ibi Treat). Microscopy was performed at room temperature with a ZEISS LSM 980 microscope with Airyscan 2, using a 63x 1.4 NA oil immersion Objective. Images were taken as z-series to generate 2D SUM projections. For determining colocalisation between Pil1-GFP and Nce102-mScarlet-I, cells were first segmented based on Nce102-mScarlet-I signal using Cellpose^108^. Segmented cells that were intersected by the image borders, and cells that featured Nce102-Scarlet signal below the fixed threshold (thresholded area = 0) for calculating Manders’ colocalization coefficient M1 (fraction of Nce102-Scarlet overlapping with Pil1-GFP) were excluded. Manders’ coefficients of single cells were obtained by analysis of 3D stacks in Fiji, using the BIOP version of the JACoP plugin^109^ with fixed manual thresholds for Pil1-GFP and Nce102-mScarlet-I, and graphs were generated with Origin Pro 2022 (OriginLab Corp.).

### Statistics and reproducibility

For MD simulations, the sample size was determined empirically, considering the time necessary for equilibration of the lipids. For our systems that is about 10 microseconds. Two replicas were performed for each lipid system. The FRAP data is combined from three individual experiments with individual measurements from these experiments pooled together for the analysis. The raw data is available upon reasonable request. For lipid sorting co-efficients, in all conditions N=2. Statistical significance was determined with Tukeýs HSD test following one-way ANOVA assuming normal distribution. Box plots elements are defined as follows: Center line is the median, box limits are 25% to 75% lower and upper quartiles, small square box inside the box limits is the mean, whiskers are the range within 1.5 IQR, circular points are datapoints, and points outside the 1.5 IQR are outliers. Yeast growth assays and microscopy were repeated at least three times on different days, yielding similar results. For calculating Manders’ overlap coefficient, microscopy data from several days was pooled to analyze at least 100 cells per mutant.

## Supporting information

Supplementary Video 1. FRAP analysis example +PI(4,5)P2/+sterol

Supplementary Video 2. FRAP analysis example -PI(4,5)P2/+sterol

Supplementary Video 3. 3DVA native eisosome lattice

## Acknowledgements

This work was supported by grants from the H2020 Marie Curie Actions IF-2020-101026765-MEMTOR (to JMK), EMBO Postdoctoral fellowships ALTF 703-2020 (to MH) and ALTF 989-2022 (to JEM), the Swiss National Supercomputing Centre (to SV) under project ID s1189 s1132 and s1221, the Swiss National Science Foundation (SNSF) under project CRSII5_189996 METEORIC (to RL, AR, SV, and JA), the European Research Council (ERC) AdG TENDO (to RL), and the SNSF under project 310030_207754 (to RL). AR and RL acknowledge additional support from the Republic and Canton of Geneva.

This work used the EM facilities at EMBL Heidelberg (iNext), the Dubochet Center for Imaging (DCI) Geneva, and DCI Lausanne, as well as the Grenoble Instruct-ERIC Center (ISBG; UAR 3518 CNRS CEA-UGA-EMBL) with support from the French Infrastructure for Integrated Structural Biology (FRISBI; ANR-10-INBS-05-02) and GRAL, a project of the University Grenoble Alpes graduate school (Ecoles Universitaires de Recherche) CBH-EUR-GS (ANR-17-EURE-0003) within the Grenoble Partnership for Structural Biology. The IBS Electron Microscope facility is supported by the Auvergne Rhône-Alpes Region, the Fonds Feder, the Fondation pour la Recherche Médicale and GIS-IBiSA.

We thank L. Tafur, M.Prouteau, A. Bergman (Loewith lab) for help with the project, M. Kaksonen and A. Picco (University of Geneva) for providing facilities and technical support for FRAP imaging and image analysis, and F. Moss III and N. Unwin for helpful discussions.

## Author contributions

J.M.K. and R.L. designed the project. J.M.K. designed experiments, processed EM data, built/refined structural models, interpreted results, and prepared manuscript. M.H. prepared reconstituted samples and designed and interpreted FRAP experiments. L.X. optimized/prepared native samples, produced CRISPR mutants. J.A. designed and interpreted MD simulations. J.E.M. designed and interpreted lipid sorting experiments. M.G.T. performed and interpreted growth assays and fluorescence microscopy. L.F.E. coded tools to assist in sorting native eisosome data. R.L. supervised the project, S.V. supervised MD simulations, A.R. supervised *in vitro* reconstitution studies, A.D. supervised EM data processing. All authors discussed the results and commented on the manuscript.

## Competing interests

The authors declare no competing interests.

## Data availability

Sharpened maps used for refinement and all associated helical maps and deepEMhancer sharpened maps have been deposited in the Electron Microscopy Data Bank under the following accession codes: Eisosome native (EMD-18307), Pil1 -PIP2/+sterol reconstituted (EMD-18308), Pil1 +PIP2/-sterol reconstituted (EMD-18309), Pil1 +PIP2/+sterol reconstituted (EMD-18310), Eisosome native compact (EMD-18311), Eisosome native stretched (EMD-18312).

Models have been deposited in the Protein Data Bank: Pil1 lattice (native) (PDB 8QB7), Lsp1 lattice (native) (PDB 8QB8), Pil1 lattice (-PIP2/+sterol reconstituted) (PDB 8QB9), Pil1 lattice (+PIP2/-sterol reconstituted) (PDB 8QBB), Pil1 lattice (+PIP2/+sterol reconstituted) (PDB 8QBD), Pil1 lattice compact (native) (PDB 8QBE), Pil1 dimer compact with lipid headgroups (native) (PDB 8QBF), and Pil1 lattice stretched (native) (PDB 8QBG).

---

**Figure Extended Data 1.**
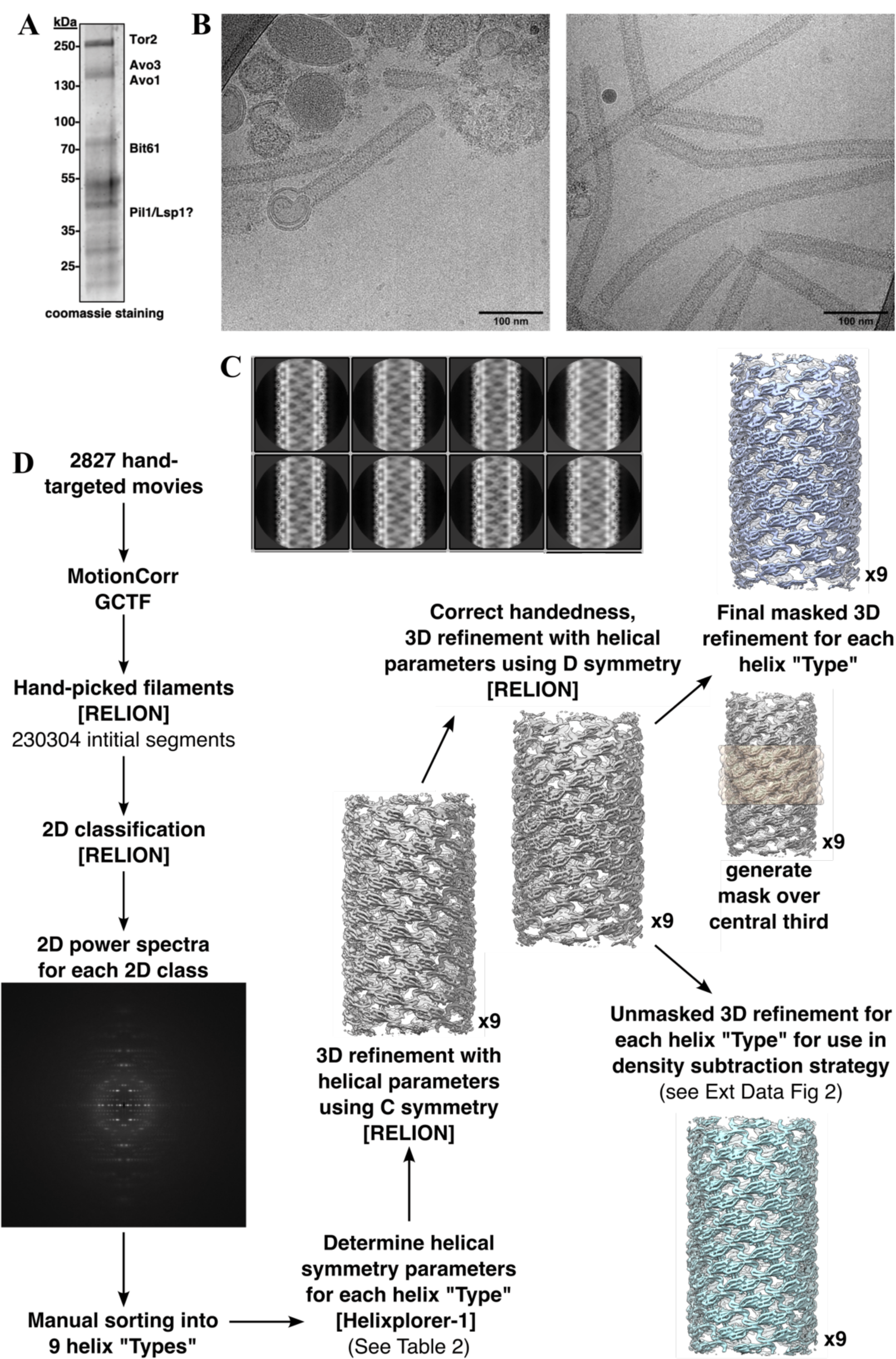
CryoEM data processing of native eisosome filaments. **A.** Coomassie staining of protein gel of Bit61-TAP purification of eisosome filaments. **B.** Raw micrographs with native eisosome filaments and other putative contaminants visible. C. Example 2D classes with varying filament diameters. **D.** Helical reconstruction data processing strategy.

**Figure Extended Data 2.**
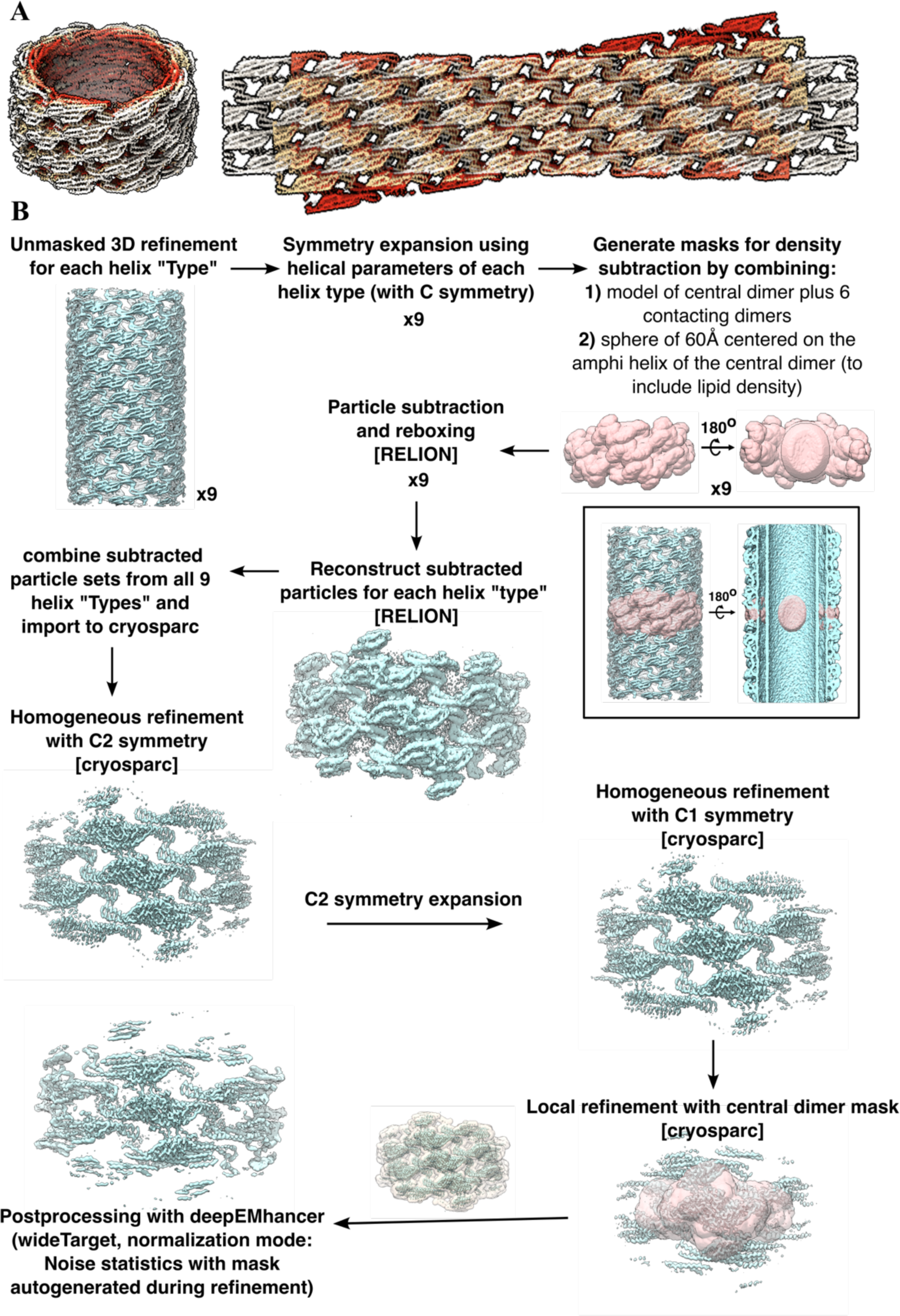
Symmetry expansion of native eisosome filaments. **A.** Unrolled and aligned helical structures of native eisosome filaments of different diameters. **B.** Data processing strategy for symmetry expansion of helical reconstructions

**Figure Extended Data 3.**
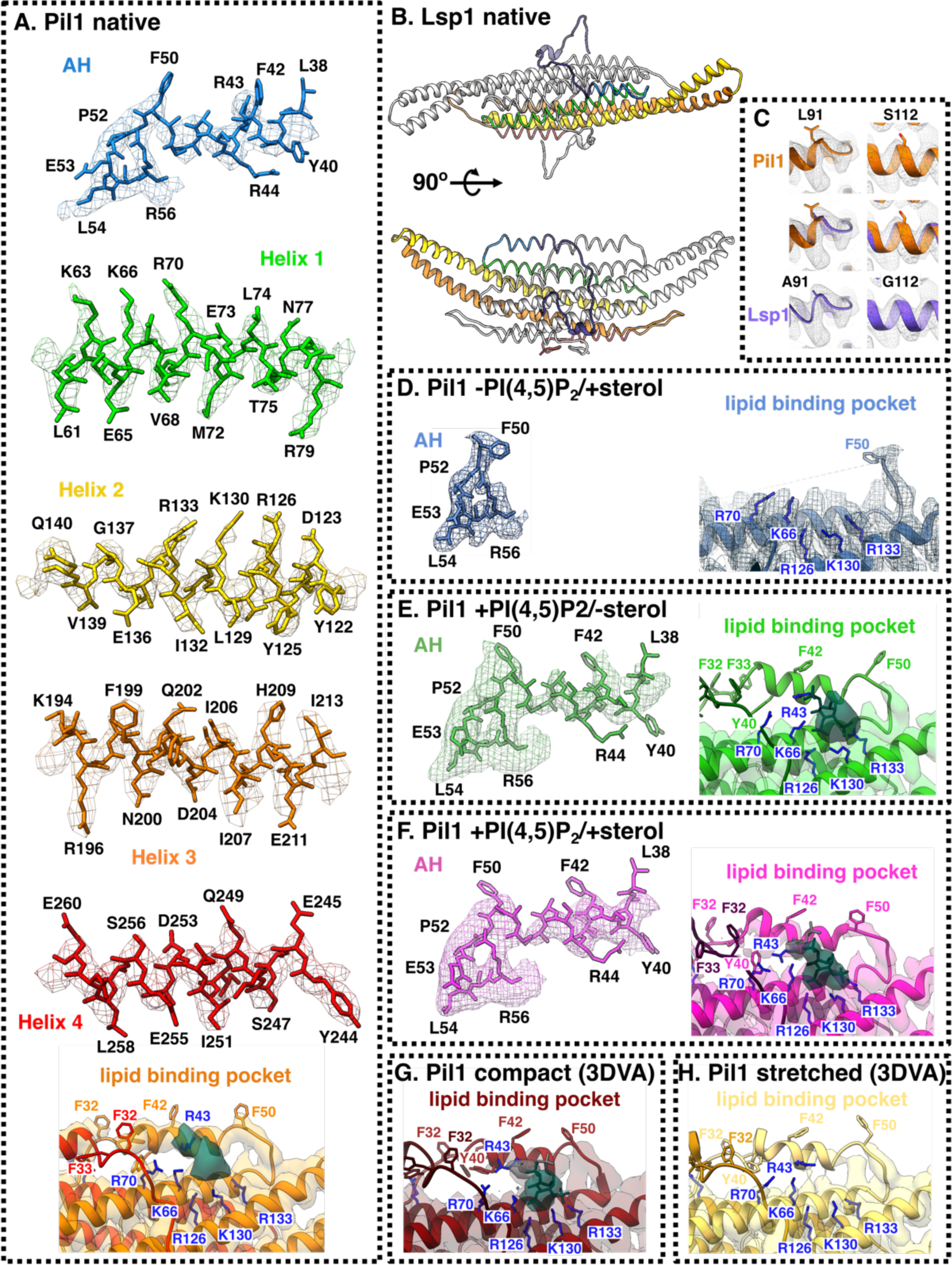
Map-to-model fit. **A.** Pil1 native eisosome map-to-model fit. **B.** Lsp1 model. **C.** Fit comparison of divergent residues in Pil1 and Lsp1. **D.** -PI(4,5)P2/+sterol reconstituted map-to-model fit of unresolved amphipathic helix and unoccupied lipid binding pocket. **E.** +PI(4,5)P2/-sterol reconstituted map-to-model fit of amphipathic helix and lipid binding pocket bound to PI(4,5)P_2_ headgroup. **F.** +PI(4,5)P2/+sterol reconstituted map-to-model fit of amphipathic helix and lipid binding pocket bound to PI(4,5)P_2_ and PS headgroups. **G.** Most compact class of 3D variability analysis (3DVA) map-to-model fit of lipid binding pocket bound to PI(4,5)P_2_ and PS headgroups. **H.** Most stretched class of 3DVA map-to-model fit of unoccupied lipid binding pocket.

**Figure Extended Data 4.**
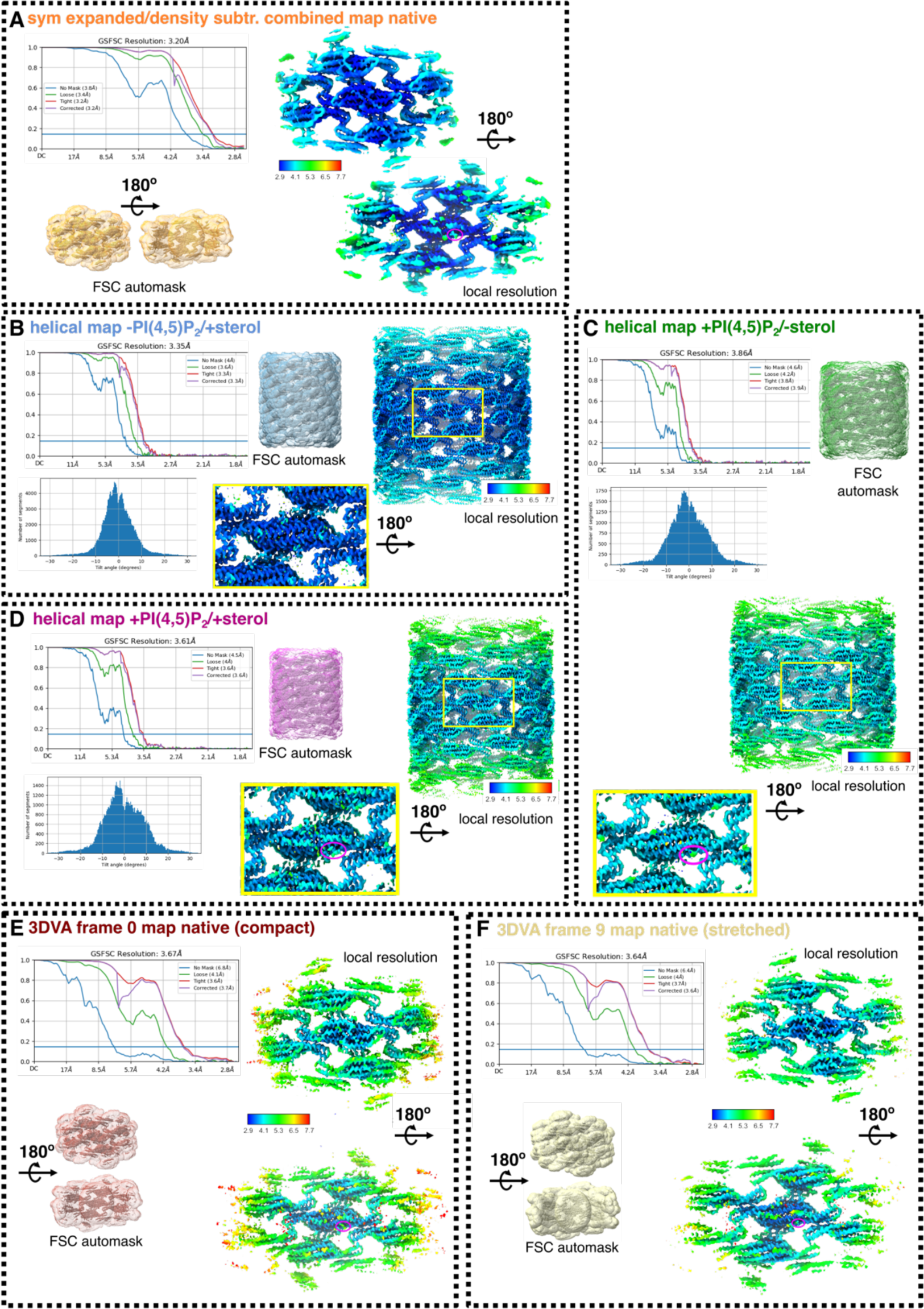
Map quality and local resolution. **A.** Gold standard Fourier shell correlation (GSFSC) plots, orientation distribution plot, auto-generated mask for average resolution determination at FSC 0.143, and local resolution map for symmetry expanded native map. Magenta circle highlights lipid binding pocket. **B-D.** GSFSC plots, helical symmetry error plot, auto-generated mask for average resolution determination at FSC 0.143, and local resolution map for helical maps of -PI(4,5)P2/+sterol (**B**), +PI(4,5)P2/-sterol (**C**), and +PI(4,5)P2/+sterol (**D**) reconstituted Pil1 tubules. Magenta circles highlight lipid binding pocket. **E-F.** GSFSC plots, orientation distribution plot, auto-generated mask for average resolution determination at FSC 0.143, and local resolution map for 3DVA frame 0 (most compact) (**E**) and 3DVA frame 9 (most stretched) (**F**) refined maps.

**Figure Extended Data 5.**
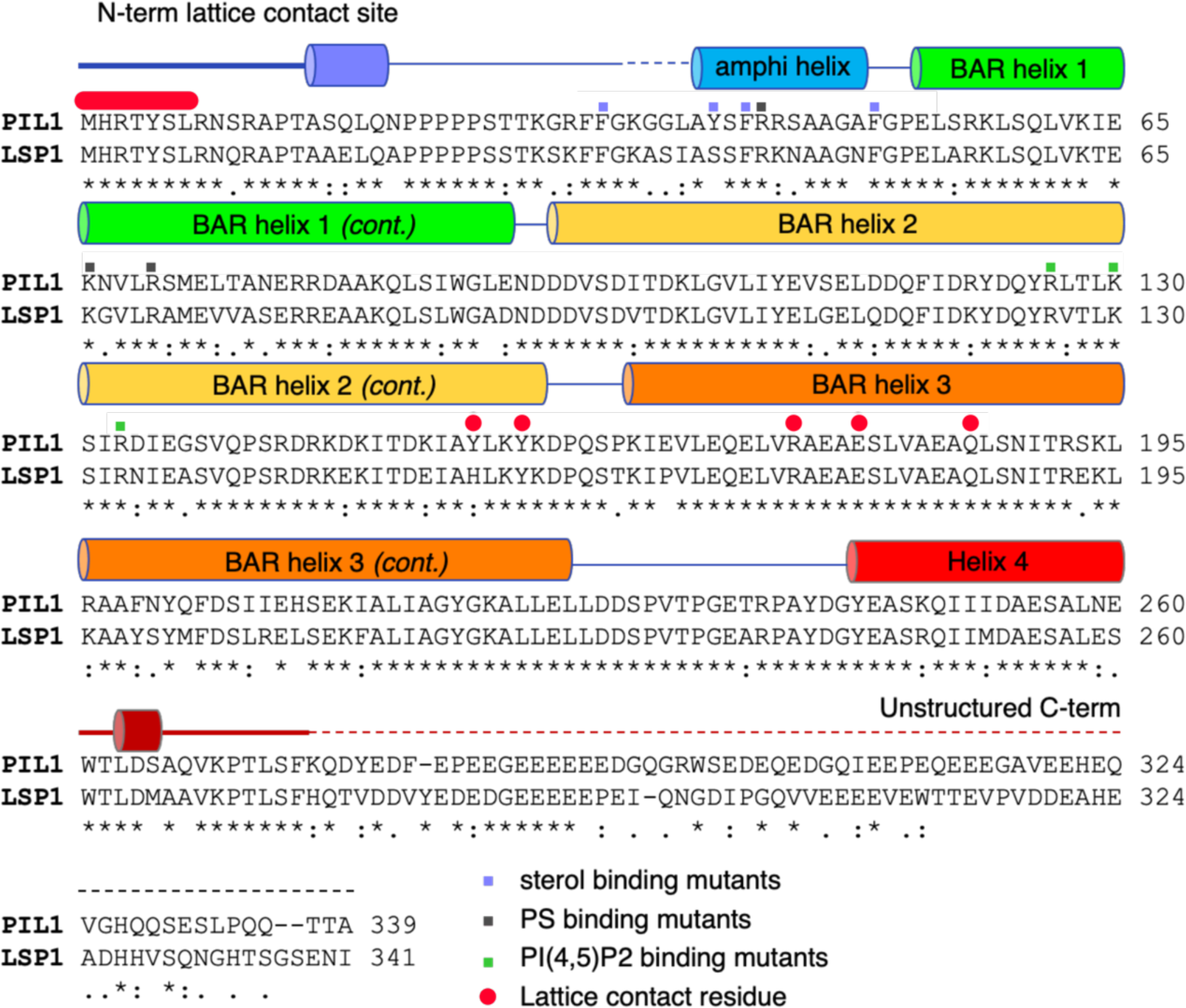
Sequence alignment of S. cerevisiae Pil1 and Lsp1. Violet squares indicate sterol binding residues, grey squares indicate proposed PS binding residues, green squares indicate PI(4,5)P_2_ binding residues, red circles indicate residues that form lattice contacts.

**Figure Extended Data 6.**
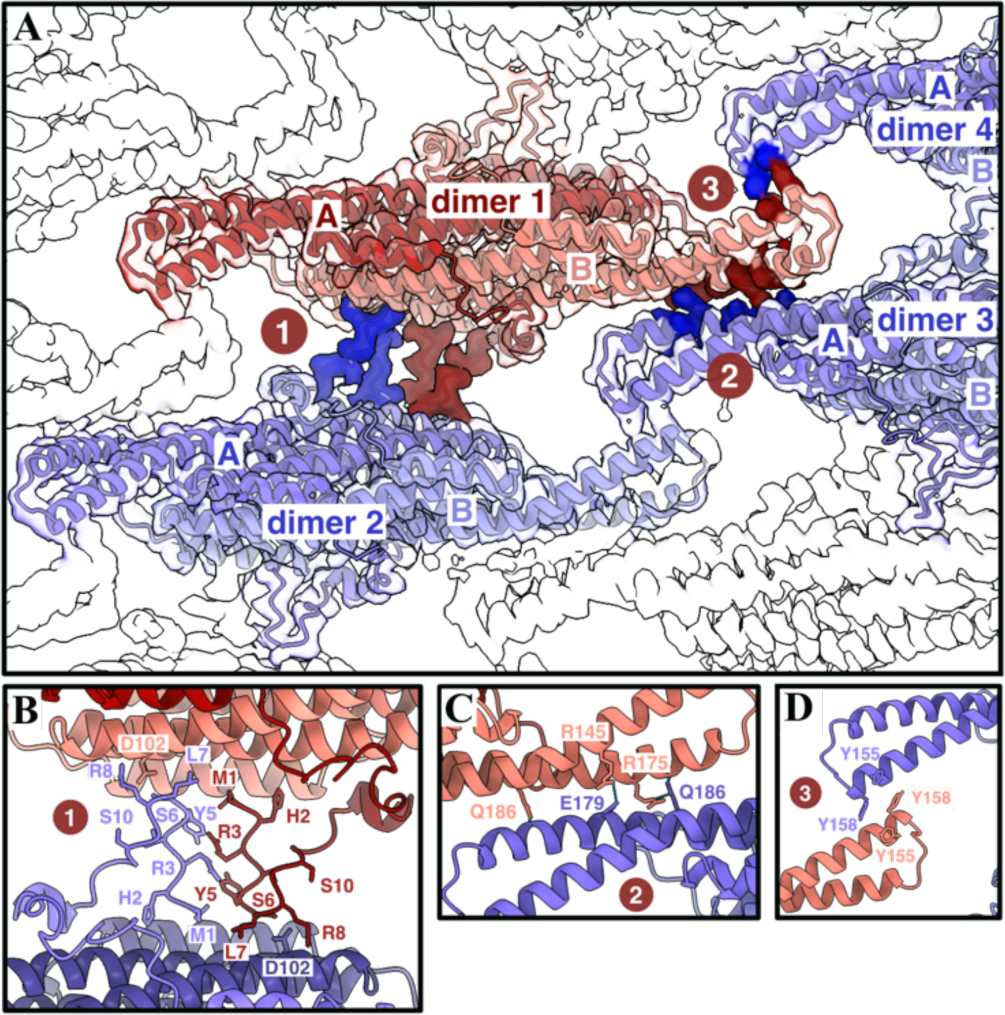
Lattice contact sites between Pil1 dimers. **A.** Overview of contact sites. **B.** Site 1: Nt contact sites formed by res 1-8. **C.** Site 2: Electrostatic interactions between res171-186 at helix 3 of BAR domain. **D.** Site 3: Hydrophobic interactions between Y155 and Y158 at BAR domain tips.

**Figure Extended Data 7.**
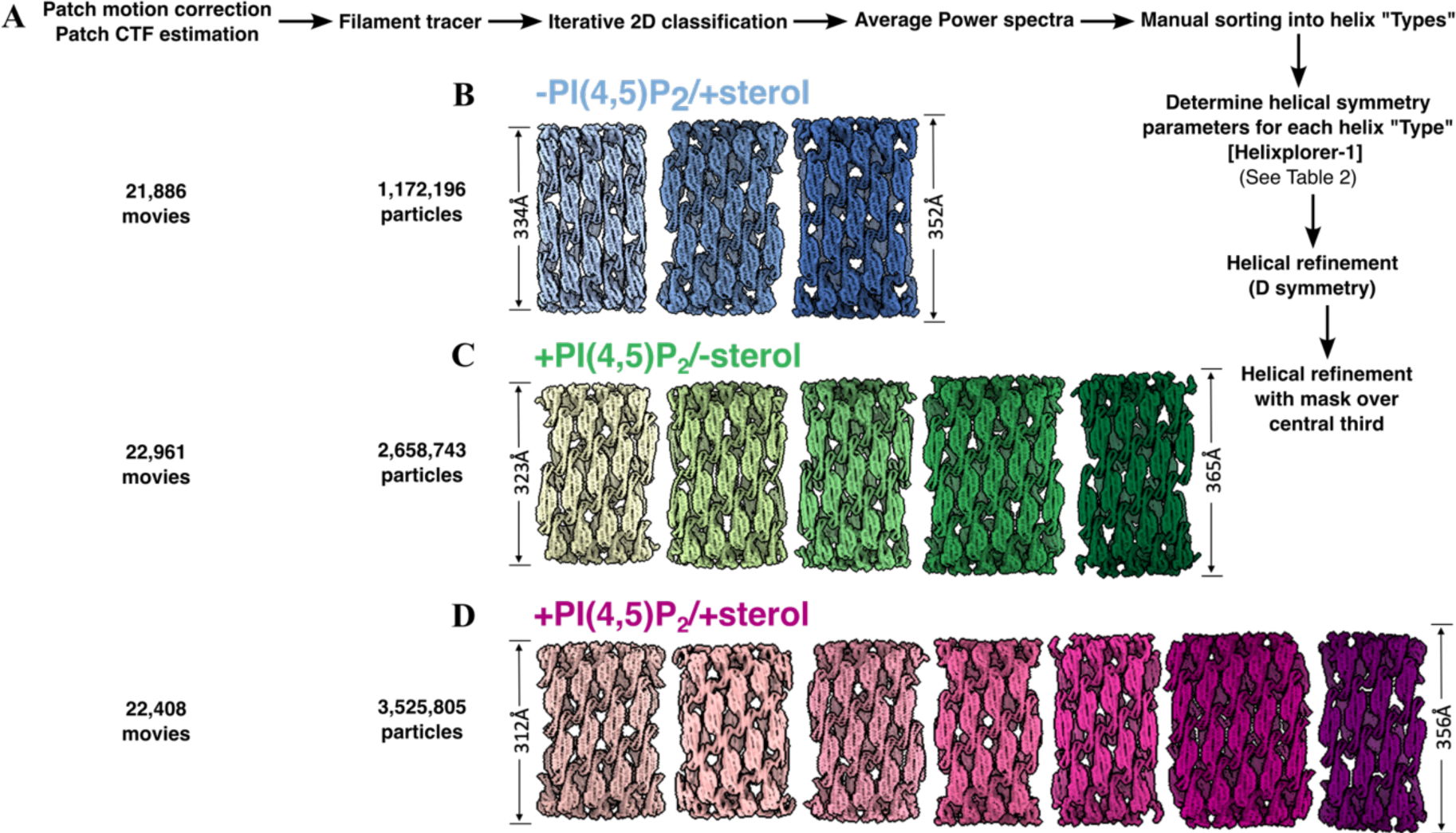
CryoEM data processing of reconstituted Pil1 filaments with lipid mixtures of known composition. **A.** Helical reconstruction data processing strategy for reconstituted Pil1 tubules. **B-D.** All helical reconstructions from -PI(4,5)P2/+sterol (**A**), +PI(4,5)P2/-sterol (**B**), and +PI(4,5)P2/+sterol (**C**) reconstituted Pil1 tubules.

**Figure Extended Data 8.**
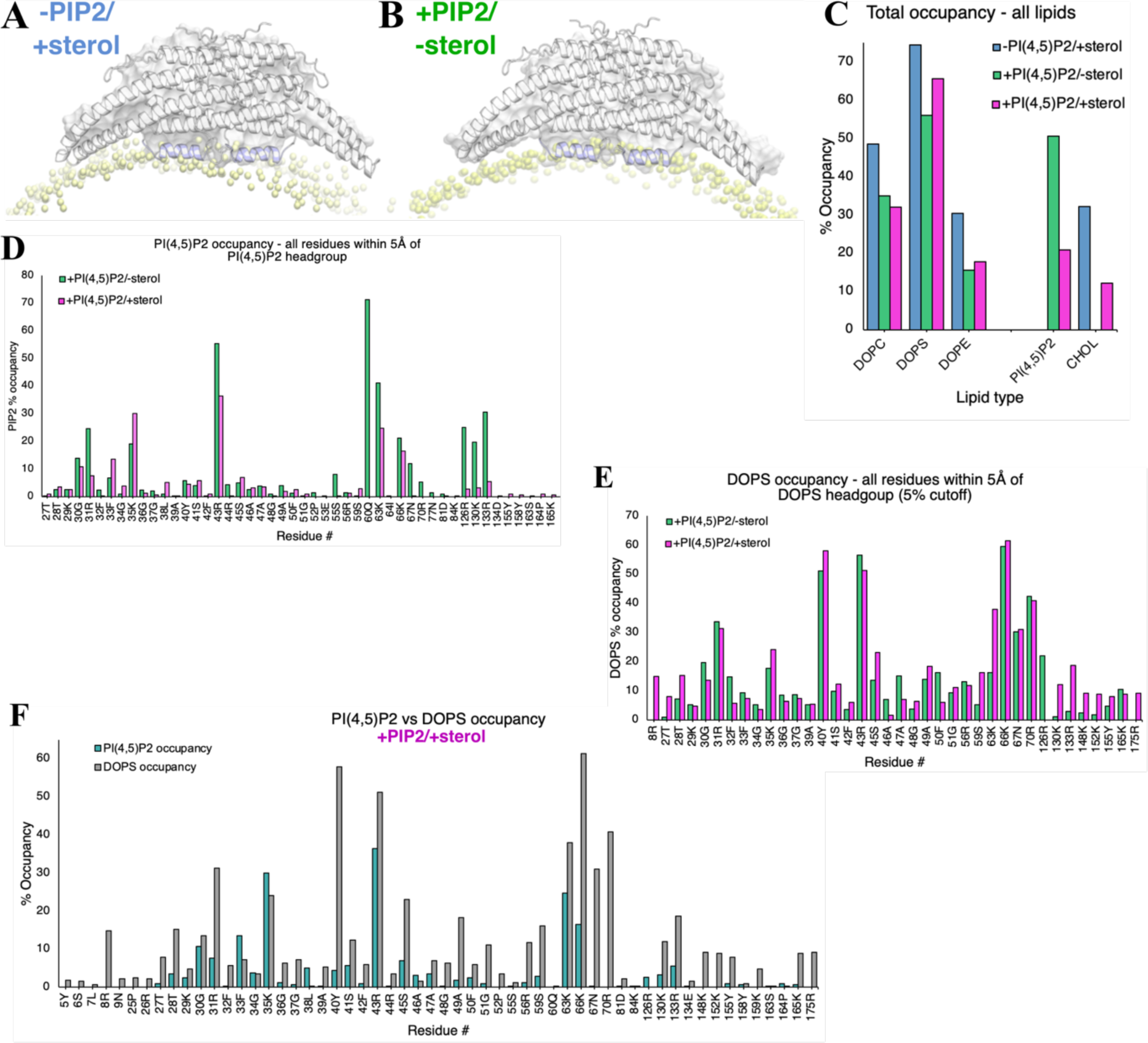
Coarse-grained molecular dynamics simulations. **A-B.** Snapshots of the amphiphilic helices (blues) in membrane (yellow) in the “-PIP2/+sterol” system (**A**) and in the +PIP2/-sterol system (**B**). **C.** Total occupancy per lipid for all lipids in each CG-MD system. **D.** PI(4,5)P_2_ occupancy reported for all residues <5Å from PI(4,5)P_2_ headgroups. **E.** DOPS lipid occupancy for residues <5Å from DOPS headgroup with >5% occupancy in CG-MD simulations. F. Comparison of PI(4,5)P_2_ occupancy and DOPS occupancy for residues <5Å from PI(4,5)P_2_ and/or DOPS headgroup.

**Figure Extended Data 9.**
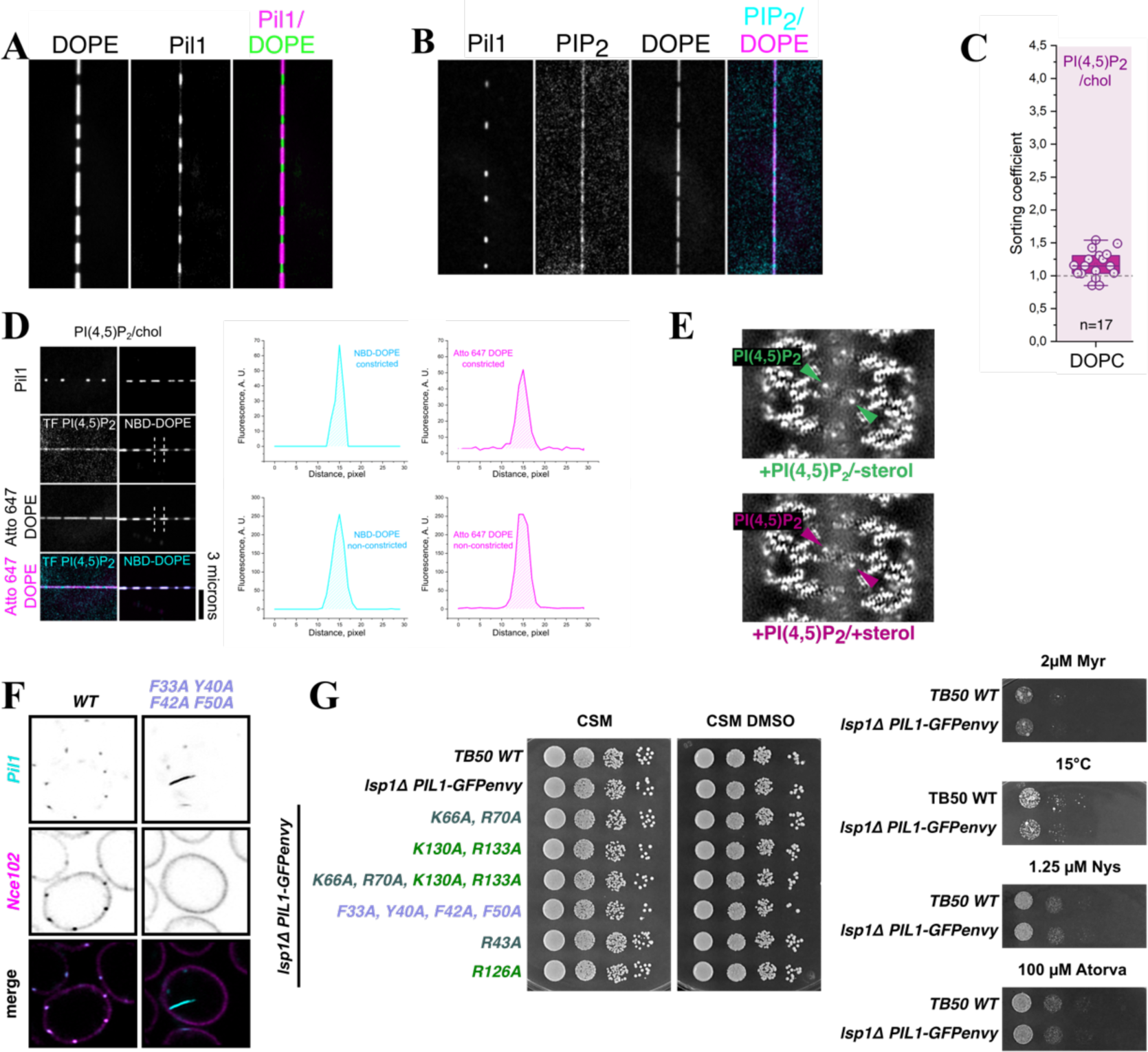
Lipid sorting co-efficients and lipid binding-impaired mutants. **A.** Example lipid nanotube with DOPE-Atto647N and Pil1-mCherry bound **B**. Example lipid nanotube with Pil1-mCherry, fluorescent TopFluor-PI(4,5)P2, and fluorescent DOPE-Atto647N. **C**. Lipid sorting co-efficient of DOPC in “+1% PI(4,5)P_2_/+sterol” reconstituted Pil1 tubules. [N=2]. **D.** Amphipathic helix slice through unsharpened maps of “+PI(4,5)P_2_/-sterol” (left panel) and “+PI(4,5)P_2_/+sterol” (right panel), with arrows indicating presumed PI(4,5)P_2_ density in each reconstruction (green and lilac, respectively). **E.** Demonstration of the fluorescence plot profiles of the lipid of interest and reference lipid used to extract the integrated fluorescence densities (hatched areas) to measure lipid sorting coefficients**. F.** Central confocal slice of *lsp1*Δ yeast expressing Pil1-GFP variants and Nce102-Scarlet-I from their endogenous loci, highlighting cytosolic ingression of eisosomes in sterol binding-impaired mutant Pil1_F33A/Y40A/F42A/F50A_. **G**. Controls for growth assays of *lsp1*Δ yeast expressing Pil1-GFP lipid binding-impaired mutants.

**Figure Extended Data 10.**
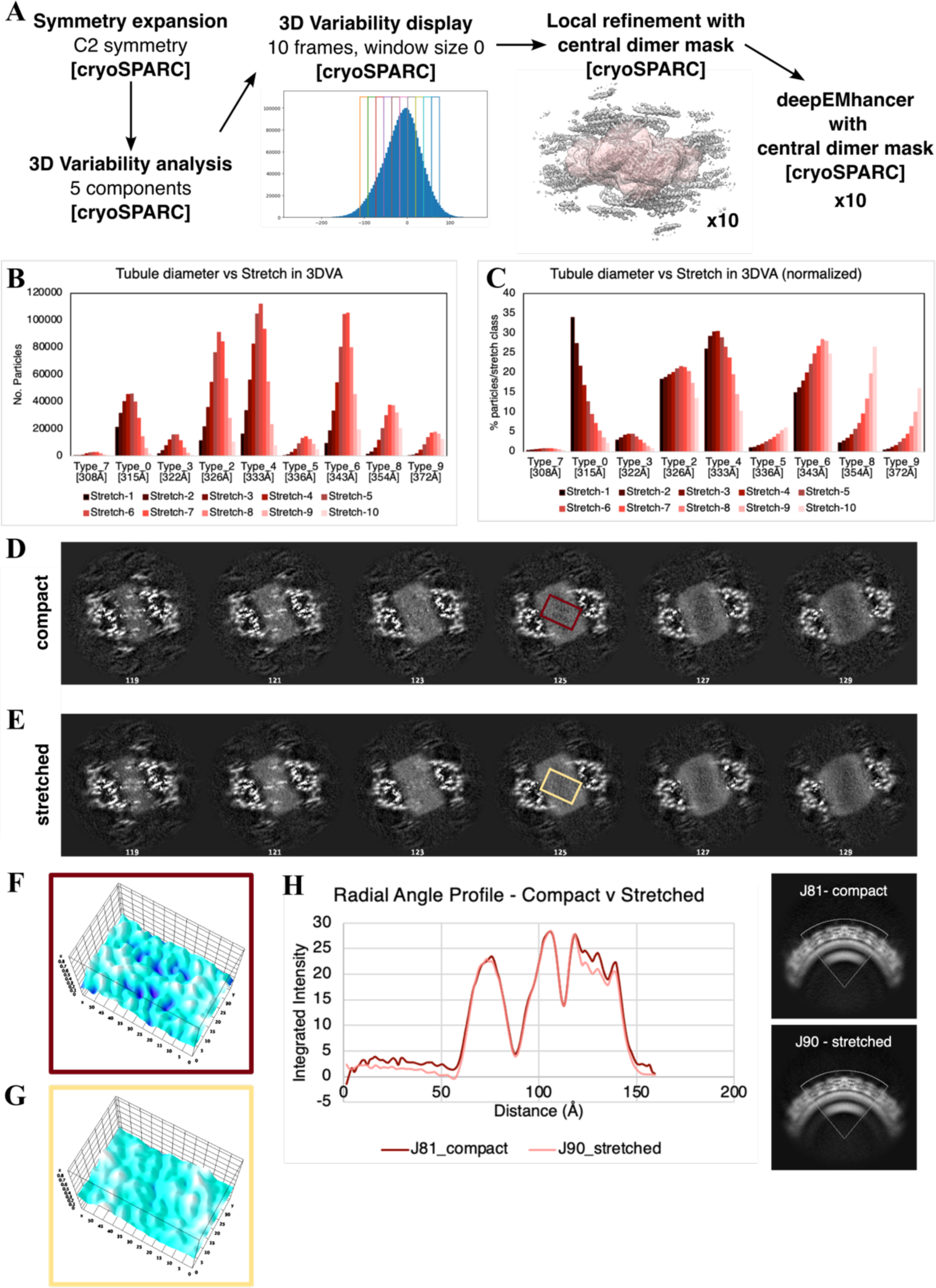
3D variability analysis. **A.** 3DVA data processing strategy. **B.** Frequency of particles from each helical reconstruction type found in each 3DVA class (1=most compact, 10=most stretched). **C.** Percent of particles normalized to total particles in each helix type found in each 3DVA class (1=most compact, 10=most stretched). **D-E**. One pixel slices separated by ∼2.6Å through unsharpened maps of most compact (**D**) and most stretched (**E**) 3DVA classes with region of maximum sterol void pattern highlighted (most compact: red box, most stretched: yellow box) **F.** 3D surface intensity plot of highlighted region (from panel D) of most compact class (red box) **G.** 3D surface intensity plot of highlighted region (from panel E) of most stretched class (yellow box). **H.** Radial angle profile plots of bilayer within most compact and most stretched classes.

**Supplementary Video 1. FRAP analysis example +PI(4,5)P_2_/+sterol.** Example FRAP of nanotube produced with +PI(4,5)P2/+sterol lipid mixture with 1% TopFluor-cholesterol (top panel, cyan) with Pil1-mCherry (middle panel, magenta) assembled on nanotube surface. Control lipid 0.1% DOPE-Atto647n shown in bottom panel (yellow).

**Supplementary Video 2. FRAP analysis example -PI(4,5)P_2_/+sterol.** Example FRAP of nanotube produced with -PI(4,5)P2/+sterol lipid mixture with 1% TopFluor-cholesterol (top panel, cyan) with Pil1-mCherry (middle panel, magenta) assembled on nanotube surface. Control lipid 0.1% DOPE-Atto647n shown in bottom panel (yellow).

**Supplementary Video 3. 3DVA native eisosome lattice.** 3D variability component from analysis using symmetry expanded/density subtracted particles from native eisosomes, demonstrating stretching at Nt lattice contact sites that expands the Pil1/Lsp1 lattice and is correlated with changes in the putative lipid-binding pocket.

**Table 1.**
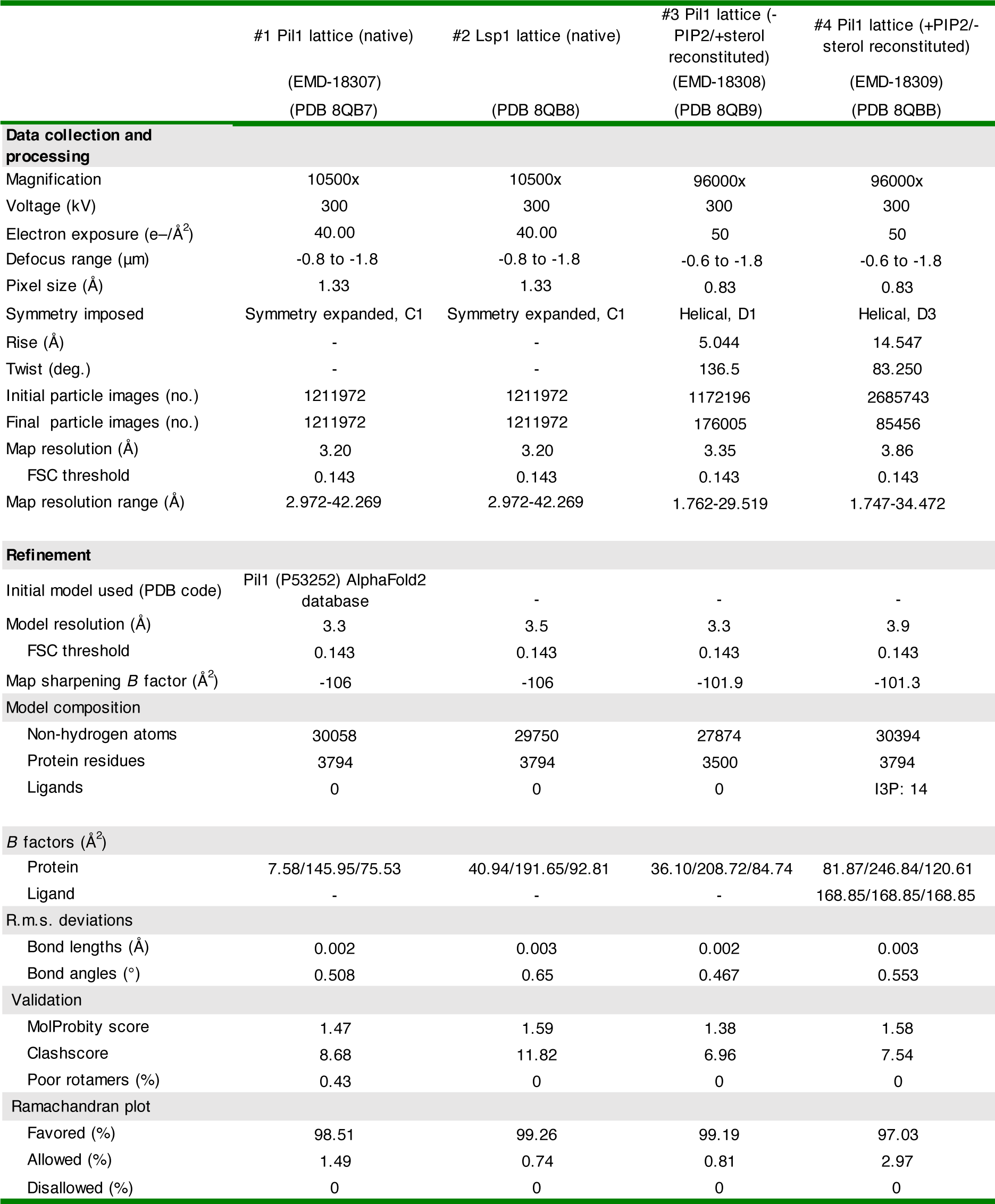
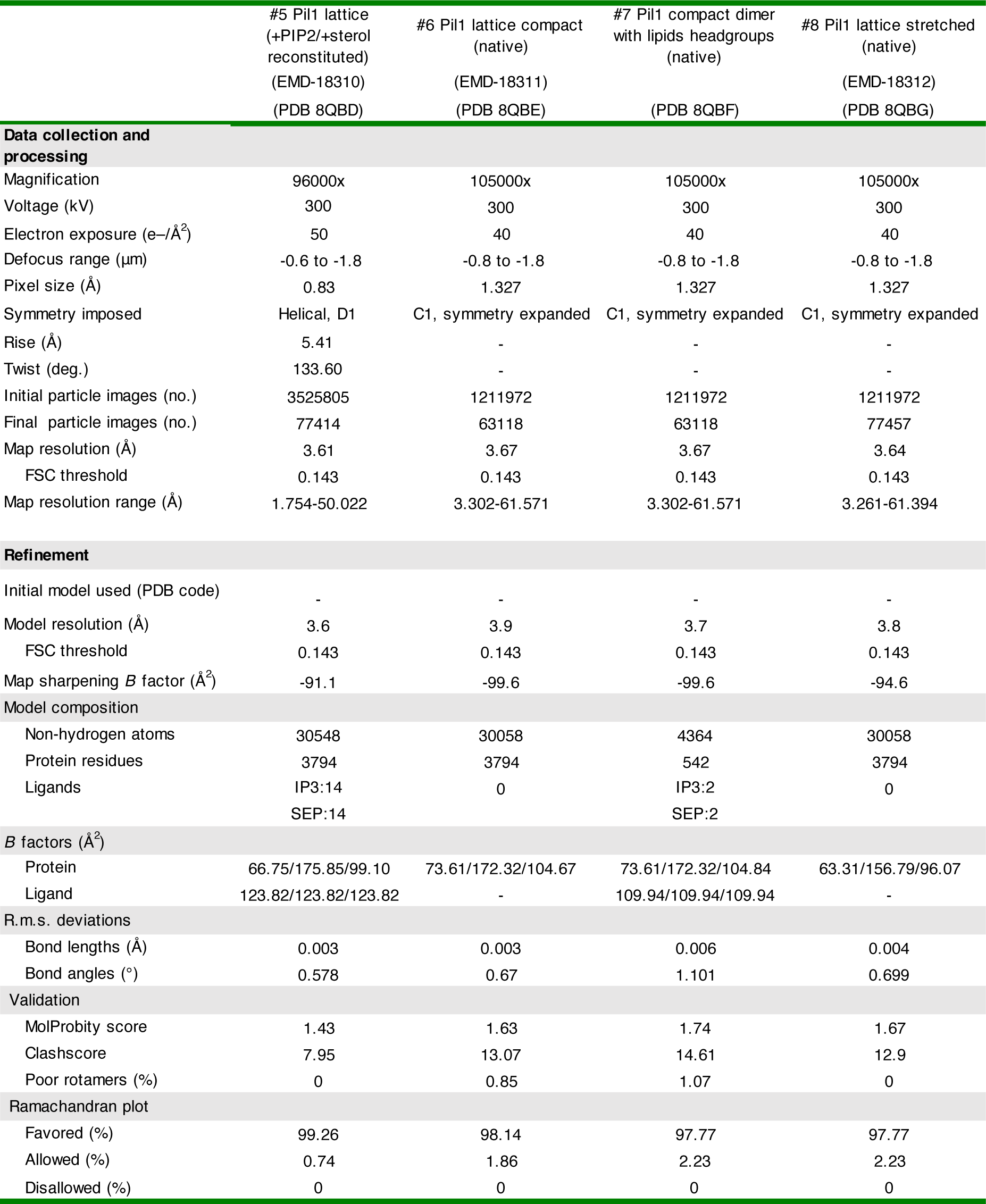
CryoEM data collection, refinement, and validation statistics.

**Table 2.**
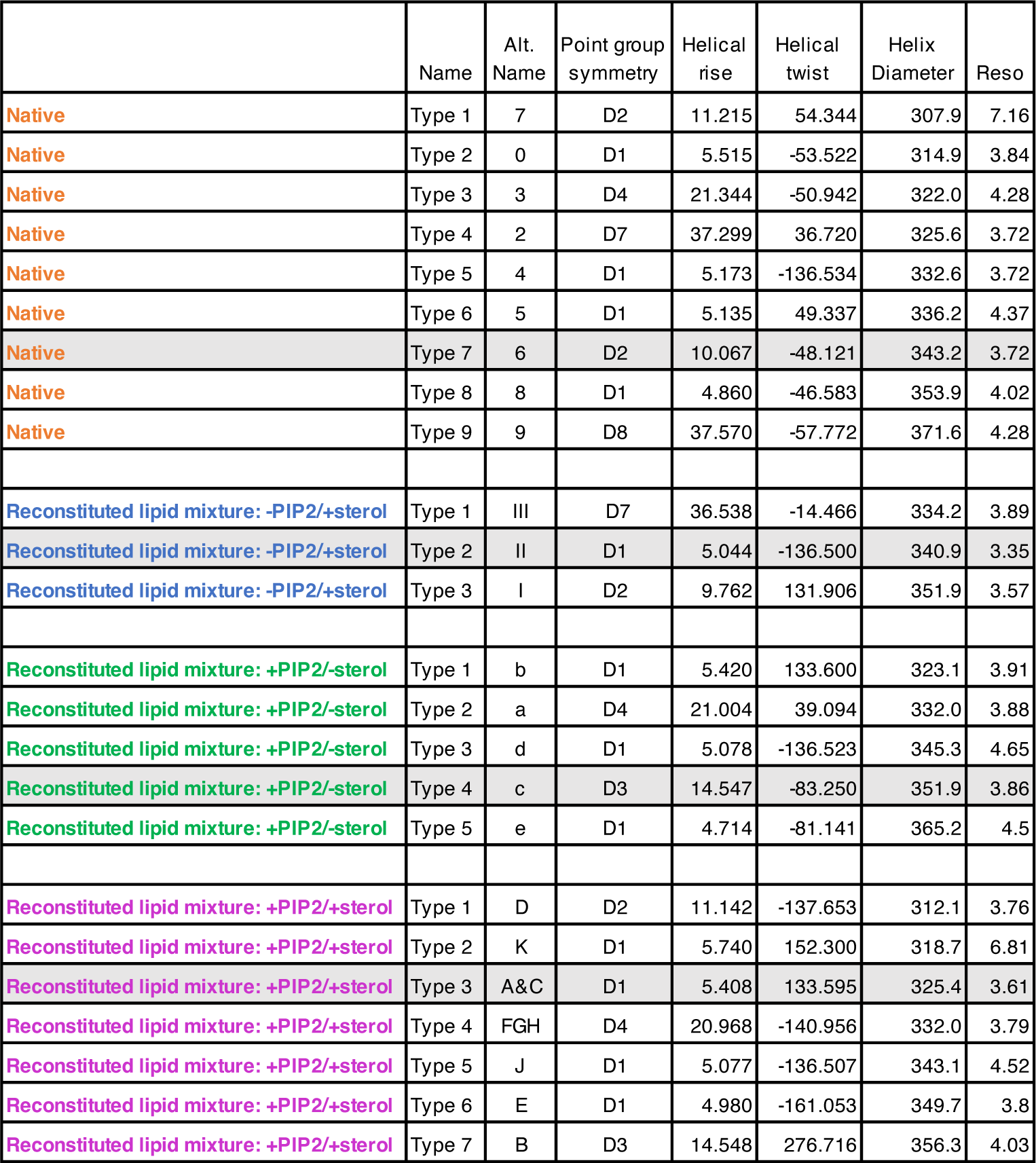
Symmetry parameters and diameters of helical structures. Helix type (numbered from smallest to largest diameter), alternative name, point group symmetry, helical rise and twist, helix diameter and resolution of helical map.

**Table 3.**
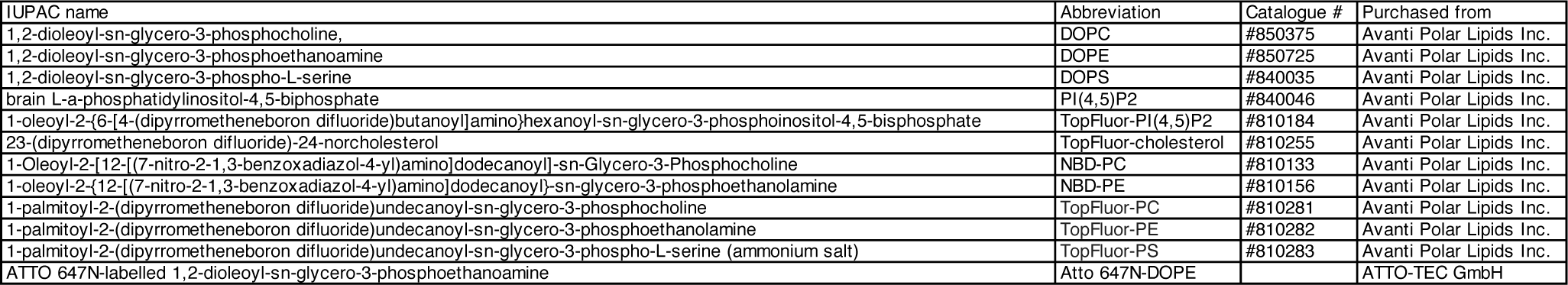
Lipids used in this study. IUPAC names, abbreviations used in text, catalogue numbers and supplier.

| IUPAC name | Abbreviation | Catalogue # | Purchased from |
| --- | --- | --- | --- |
| 1,2-dioleoyl-sn-glycero-3-phosphocholine, | DOPC | #850375 | Avanti Polar Lipids Inc. |
| 1,2-dioleoyl-sn-glycero-3-phosphoethanolamine | DOPE | #850725 | Avanti Polar Lipids Inc. |
| 1,2-dioleoyl-sn-glycero-3-phospho-L-serine | DOPS | #840035 | Avanti Polar Lipids Inc. |
| brain L-α-phosphatidylinositol-4,5-bisphosphate | PI(4,5)P2 | #840046 | Avanti Polar Lipids Inc. |
| 1-oleoyl-2-(6-[4-(dipyrrometheneboron difluoride)butanoyl]amino)hexanoyl-sn-glycero-3-phosphoinositol-4,5-bisphosphate | TopFluor-PI(4,5)P2 | #810184 | Avanti Polar Lipids Inc. |
| 23-(dipyrrometheneboron difluoride)-24-norcholesterol | TopFluor-cholesterol | #810255 | Avanti Polar Lipids Inc. |
| 1-Oleoyl-2-[12-[(7-nitro-2-1,3-benzoxadiazol-4-yl)amino]dodecanoyl]-sn-Glycero-3-Phosphocholine | NBD-PC | #810133 | Avanti Polar Lipids Inc. |
| 1-oleoyl-2-[12-[(7-nitro-2-1,3-benzoxadiazol-4-yl)amino]dodecanoyl]-sn-glycero-3-phosphoethanolamine | NBD-PE | #810156 | Avanti Polar Lipids Inc. |
| 1-palmitoyl-2-(dipyrrometheneboron difluoride)undecanoyl-sn-glycero-3-phosphocholine | TopFluor-PC | #810281 | Avanti Polar Lipids Inc. |
| 1-palmitoyl-2-(dipyrrometheneboron difluoride)undecanoyl-sn-glycero-3-phosphoethanolamine | TopFluor-PE | #810282 | Avanti Polar Lipids Inc. |
| 1-palmitoyl-2-(dipyrrometheneboron difluoride)undecanoyl-sn-glycero-3-phospho-L-serine (ammonium salt) | TopFluor-PS | #810283 | Avanti Polar Lipids Inc. |
| ATTO 647N-labelled 1,2-dioleoyl-sn-glycero-3-phosphoethanolamine | Atto 647N-DOPE |  | ATTO-TEC GmbH |

**Table 4.** Lipid mixtures used in this study. Mixture name used in text, components of lipid mixtures, and molar ratio (mol %).

**CryoEM studies**
| Name | Lipid mixture | molar ratio |
| --- | --- | --- |
| -PI(4,5)P2/+sterol | DOPC:DOPE:DOPS:Cholesterol | 30:20:20:30 mol% |
| +PI(4,5)P2/-sterol | DOPC:DOPE:DOPS:brain PI(4,5)P2 | 50:20:20:10 mol% |
| +PI(4,5)P2/+sterol | DOPC:DOPE:DOPS:Cholesterol:brain PI(4,5)P2 | 35:20:20:15:10 mol% |

**Lipid sorting coefficient and FRAP studies\***
| Name | Lipid mixture | molar ratio |
| --- | --- | --- |
| -PI(4,5)P2/+sterol | DOPC:DOPE:DOPS:cholesterol:TopFluor-cholesterol | 30:20:20:29:1 mol% |
| +1% PI(4,5)P2/+sterol | DOPC:DOPE:DOPS:cholesterol:TopFluor-cholesterol:brain PI(4,5)P2 | 29:20:20:29:1:1 mol% |
| +1% PI(4,5)P2/-sterol | DOPC:DOPE:DOPS:TopFluor-PI(4,5)P2 | 59:20:20:1 mol% |
| +1% PI(4,5)P2/+sterol | DOPC:DOPE:DOPS:cholesterol:TopFluor-PI(4,5)P2 | 29:20:20:30:1 mol% |
| +1% PI(4,5)P2/+sterol | DOPC:DOPE:DOPS:cholesterol:brain P(4,5)P2:TopFluor-PC | 28:20:20:30:1:1 mol% |
| +1% PI(4,5)P2/+sterol | DOPC:DOPE:DOPS:cholesterol:brain P(4,5)P2:TopFluor-PE | 29:19:20:30:1:1 mol% |
| +1% PI(4,5)P2/+sterol | DOPC:DOPE:DOPS:cholesterol:brain P(4,5)P2:TopFluor-PS | 29:20:19:30:1:1 mol% |
| +1% PI(4,5)P2/+sterol | DOPC:DOPE:DOPS:cholesterol:brain P(4,5)P2:NBD-PE | 29:19:20:30:1:1 mol% |
| +1% PI(4,5)P2/+sterol | DOPC:DOPE:DOPS:cholesterol:brain P(4,5)P2:NBD-PC | 29:19:20:30:1:1 mol% |

\*0.01 mol% of Atto647N DOPE was added to each lipid mixture for these experiments

**Table 5.** FRAP halftimes in seconds and percentages of mobile fractions. Values are derived from one-phase exponential (one halftime shown) or two-phase exponential (two halftimes shown) equations (see Materials and methods). Standard error of fitted values (s.e.) and the goodness of fits (R-squared) are shown.

**TopFluor-cholesterol**
|  | halftime 1 (s.e.), s | halftime 2 (s.e.), s | mobile fraction (s.e.), % | R-squared |
| --- | --- | --- | --- | --- |
| +PIP2, control | 5.6 (0.4) | 64.1 (8.6) | 86.4 (0.5) | 0.993 |
| +PIP2, Pil1 | 8.7 (0.3) | - | 34.0 (0.1) | 0.953 |
| -PIP2, control | 5.0 (1.0) | 38.5 (3.6) | 92.5 (0.9) | 0.987 |
| -PIP2, Pil1 | 6.7 (0.2) | - | 56.3 (0.1) | 0.971 |

**TopFluor-PIP2**
|  | halftime 1 (s.e.), s | halftime 2 (s.e.), s | mobile fraction (s.e.), % | R-squared |
| --- | --- | --- | --- | --- |
| +sterol, control | 4.8 (0.3) | 35.4 (1.2) | 60.8 (0.4) | 0.998 |
| +sterol, Pil1 | 4.6 (0.6) | 61.9 (4.4) | 20.7 (0.6) | 0.994 |
| -sterol, control | 4.6 (0.3) | 27.3 (2.0) | 56.7 (0.4) | 0.994 |
| -sterol, Pil1 | 28.5 (1.0) | - | 27.5 (0.3) | 0.973 |

**TopFluor-PC**
|  | halftime 1 (s.e.), s | halftime 2 (s.e.), s | mobile fraction (s.e.), % | R-squared |
| --- | --- | --- | --- | --- |
| +PIP2, control | 1.3 (0.4) | 12.3 (0.4) | 75.1 (0.3) | 0.991 |
| +PIP2, Pil1 | 14.4 (0.4) | - | 58.7 (0.3) | 0.996 |
| -PIP2, control | 2.7 (0.3) | 18.8 (0.5) | 90.7 (0.5) | 0.998 |
| -PIP2, Pil1 | 1.0 (0.1) | 13.2 (0.4) | 55.9 (0.2) | 0.968 |

**TopFluor-PE**
|  | halftime 1 (s.e.), s | halftime 2 (s.e.), s | mobile fraction (s.e.), % | R-squared |
| --- | --- | --- | --- | --- |
| +PIP2, control | 9.7 (0.1) | - | 80.4 (0.2) | 0.988 |
| +PIP2, Pil1 | 9.0 (0.3) | - | 49.9 (0.2) | 0.926 |
| -PIP2, control | 1.9 (0.3) | 15.6 (0.3) | 81.6 (0.2) | 0.995 |
| -PIP2, Pil1 | 0.8 (0.1) | 7.6 (0.5) | 54.4 (0.1) | 0.951 |

**TopFluor-PS**
|  | halftime 1 (s.e.), s | halftime 2 (s.e.), s | mobile fraction (s.e.), % | R-squared |
| --- | --- | --- | --- | --- |
| +PIP2, control | 2.9 (0.2) | 21.7 (0.5) | 86.5 (0.2) | 0.997 |
| +PIP2, Pil1 | 12.7 (0.2) | - | 55.4 (0.2) | 0.982 |
| -PIP2, control | 2.6 (0.2) | 17.7 (0.4) | 73.0 (0.1) | 0.997 |
| -PIP2, Pil1 | 0.7 (0.2) | 7.8 (0.4) | 49.6 (0.1) | 0.954 |

**Table 6.**
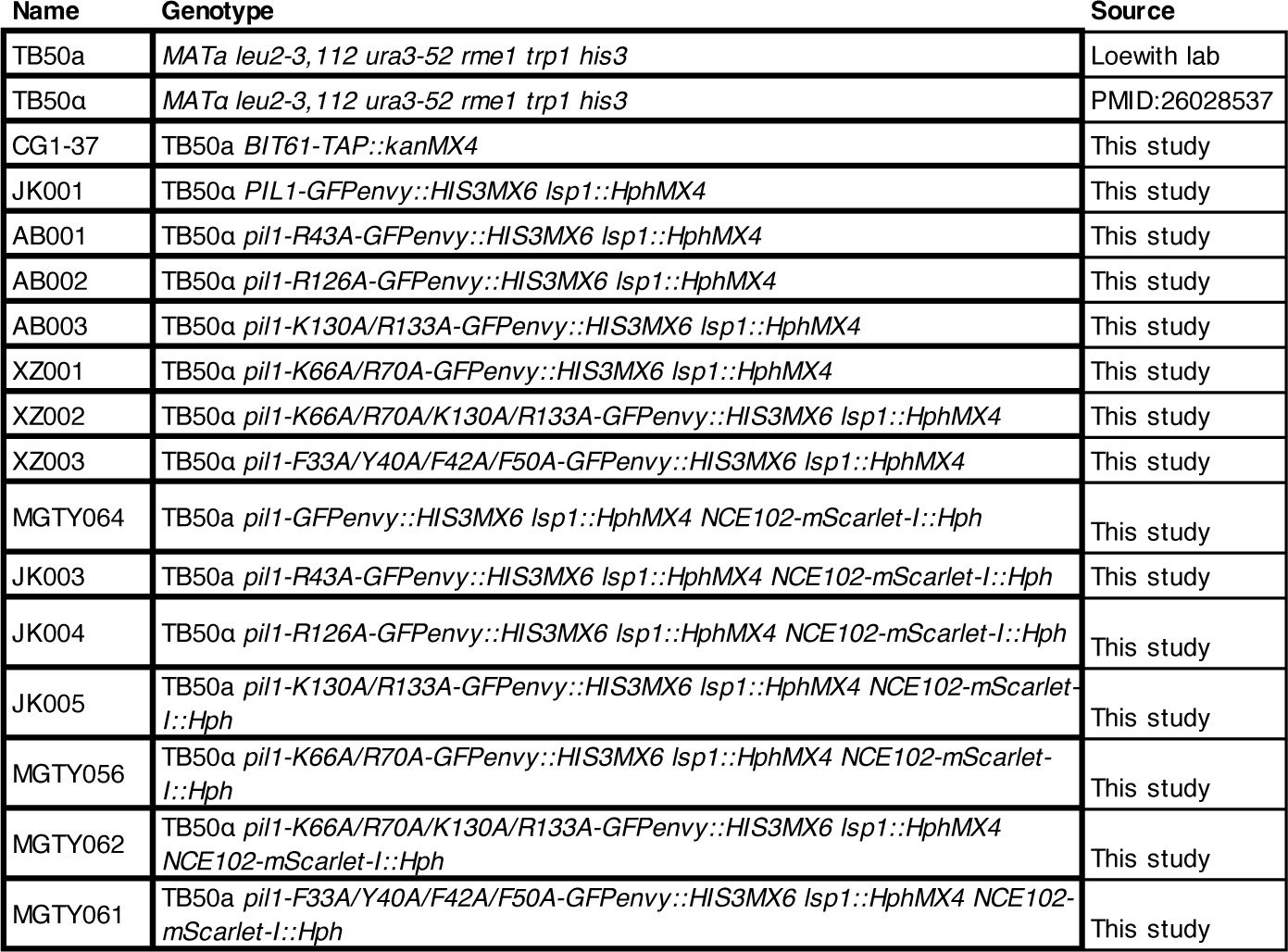
Yeast strains used in this study. Strain name, genotype, and source.

## Notes

### Competing Interest Statement

The authors have declared no competing interest.

